# The *Kobresia pygmaea* ecosystem of the Tibetan highlands – origin, functioning and degradation of the world’s largest pastoral alpine ecosystem

**DOI:** 10.1101/135558

**Authors:** Georg Miehe, Per-Marten Schleuss, Elke Seeber, Wolfgang Babel, Tobias Biermann, Martin Braendle, Fahu Chen, Heinz Coners, Thomas Foken, Tobias Gerken, Hans-F. Graf, Georg Guggenberger, Silke Hafner, Maika Holzapfel, Johannes Ingrisch, Yakov Kuzyakov, Zhongping Lai, Lukas Lehnert, Christoph Leuschner, Jianquan Liu, Shibin Liu, Yaoming Ma, Sabine Miehe, Volker Mosbrugger, Henry J. Noltie, Lars Opgenoorth, Joachim Schmidt, Sandra Spielvogel, Sebastian Unteregelsbacher, Yun Wang, Sandra Willinghöfer, Xingliang Xu, Yongping Yang, Shuren Zhang, Karsten Wesche

## Abstract

*Kobresia* pastures in the eastern Tibetan highlands occupy 450000 km^2^ and form the world’s largest pastoral alpine ecosystem. The main constituent is an endemic dwarf sedge, *Kobresia pygmaea*, which forms a lawn with a durable turf cover anchored by a felty root mat, and occurs from 3000 m to nearly 6000 m a.s.l. The existence and functioning of this unique ecosystem and its turf cover have not yet been explained against a backdrop of natural and anthropogenic factors, and thus its origin, drivers, vulnerability or resilience remain largely unknown. Here we present a review on ecosystem diversity, reproduction and ecology of the key species, pasture health, cycles of carbon (C), water and nutrients, and on the paleo-environment. The methods employed include molecular analysis, grazing exclusion, measurements with micro-lysimeters and gas exchange chambers, ^13^C and ^15^N labelling, eddy-covariance flux measurements, remote sensing and atmospheric modelling.

The following combination of traits makes *Kobresia pygmaea* resilient and highly competitive: dwarf habit, predominantly below-ground allocation of photo assimilates, mixed reproduction strategy with both seed production and clonal growth, and high genetic diversity. Growth of *Kobresia* pastures is co-limited by low rainfall during the short growing season and livestock-mediated nutrient withdrawal. Overstocking has caused pasture degradation and soil deterioration, yet the extent remains debated. In addition, we newly describe natural autocyclic processes of turf erosion initiated through polygonal cracking of the turf cover, and accelerated by soil-dwelling endemic small mammals. The major consequences of the deterioration of the vegetation cover and its turf include: (1) the release of large amounts of C and nutrients and (2) earlier diurnal formation of clouds resulting in (3) decreased surface temperatures with (4) likely consequences for atmospheric circulation on large regional and, possibly global, scales.

Paleo-environmental reconstruction, in conjunction with grazing experiments, suggests that the present grazing lawns of *Kobresia pygmaea* are synanthropic and may have existed since the onset of pastoralism. The traditional migratory rangeland management was sustainable over millennia and possibly still offers the best strategy to conserve, and possibly increase, the C stocks in the *Kobresia* turf, as well as its importance for climate regulation.

## INTRODUCTION

The Tibetan highlands encompass 90% of the Earth’s terrain above 4000 m and host the world’s largest pastoral alpine ecosystem: the *Kobresia* pastures of the south-eastern highlands, with an area of 450000 km^2^ (Fig. 1). This ecosystem is globally unique as it is:

**FIG. 1.**
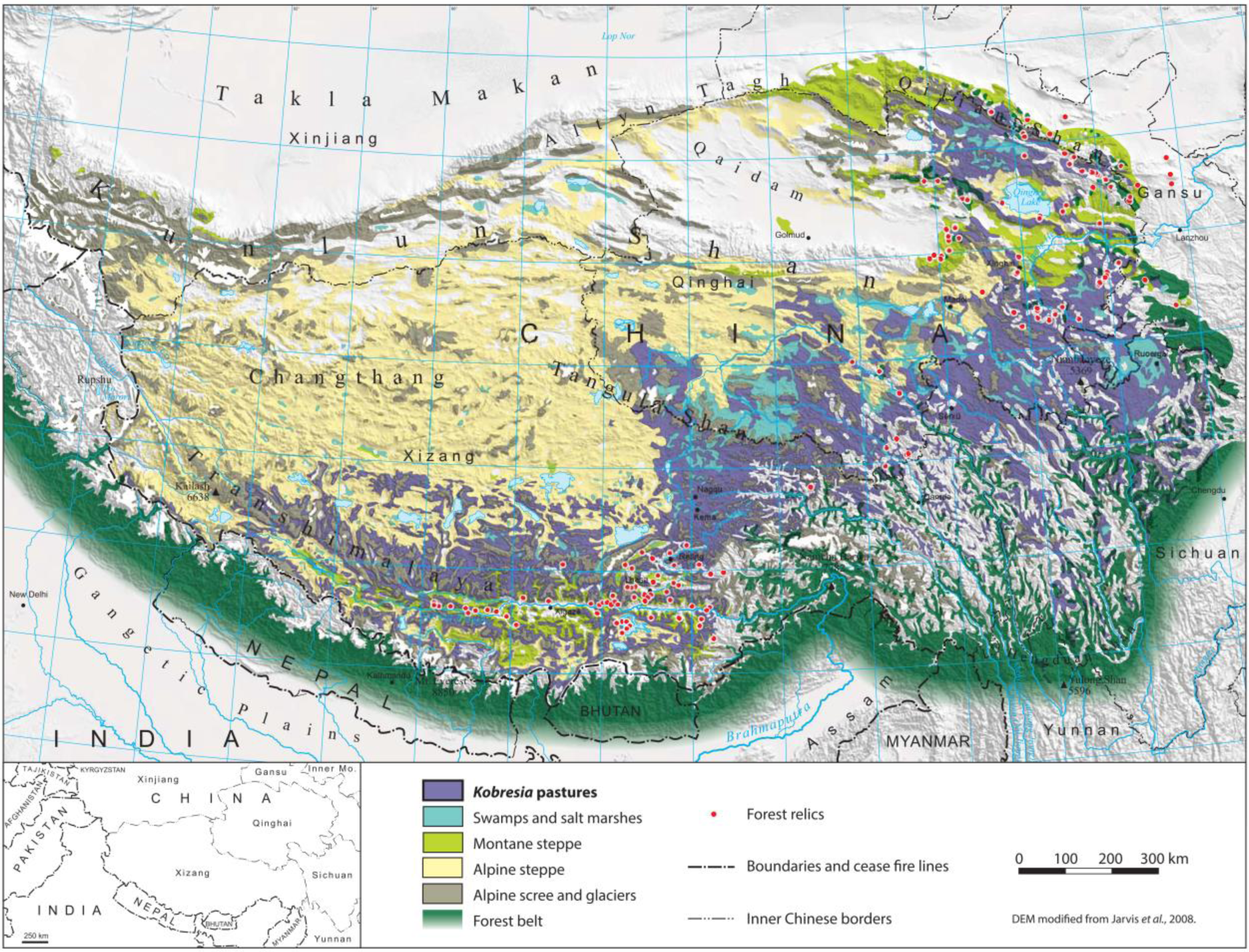
*Kobresia pygmaea* pastures of the Tibetan highlands and forest relics. After Atlas Tibet Plateau 1990, Miehe et al. 2008*b*, 2014, Babel et al. 2014.

1. dominated by a single endemic sedge species of 1 to 4 cm in height;
2. forms a golf-course like lawn, with a very durable turf cover anchored by a felty root mat;
3. occurs over an elevational extent of 3000 m, stretching between 3000 m (in the north-eastern highlands) to nearly 6000 m (on the north slope of Mt. Everest; Miehe 1989, Miehe *et al.* 2008*b*).

The evolution and recent changes of the *Kobresia* ecosystem, as well as its future development, are of great importance because surface properties of the highlands have an undisputed effect on the global climate (Cui and Graf 2009, Babel et al. 2014, Yang et al. 2014). The Tibetan plateau is significant in terms of carbon (C) turnover and CO_2_ fluxes from regional to global scales (Zhao et al. 2005, Zhang et al. 2007). Its soil stores huge C amounts, mounting up to 2.5% to the global C pool (Ni 2002, Wang et al. 2002, Hafner et al. 2012). Of these, the *Kobresia* ecosystem in particular contributes about 1% (Batjes 1996, Wang et al. 2002), though covering only about 0.3% of the global land surface (equivalent to one-quarter of the whole Tibetan plateau). The livelihoods of 5 million pastoralists depend on forage resources from the rangelands, which sustain about 13 million yak, and 30 million goats and sheep (Wiener et al. 2003, Suttie et al. 2005). One quarter of the world’s population living in the surrounding lowlands ultimately depend on ecosystem functions of the *Kobresia* mats, which represent the upper catchment areas of the Huang He, Yangtze, Salween, Mekong, and partly of the Brahmaputra, rivers.

The aim of this review is therefore to summarize recent findings relating to:

1. the diversity and the distribution of plant species of the ecosystem and its paleo-ecological background;
2. the ecology and reproduction of the dominant species, *Kobresia pygmaea* (C.B.Clarke) C.B.Clarke;
3. the ecosystem’s water budget and hydrological fluxes;
4. fluxes in the carbon cycle;
5. soil properties and functions, including productivity and nutrients stocks;
6. the main causes of current rangeland degradation and its consequences;
7. the human impact shaping this ecosystem;
8. the current understanding of the age of this human impact.

In addition, a new concept of natural autocyclic processes of turf erosion, initiated through polygonal cracking of the turf cover increased by overgrazing and facilitated by soil-dwelling endemic small mammals, will be presented.

Building on available literature, this review integrates field observations from surveys in the highlands undertaken by the first author between 1984 and 2016, and related studies on the status and dynamics of the ecosystem from grazing exclusion-experiments, obtained mainly in the northeastern montane *Kobresia* rangelands near Xinghai (Qinghai Province, 3440 m, 35°32’N / 99°51’E), and in the core area of alpine *Kobresia pygmaea* pastures next to the ‘*Kobresia pygmaea* Research Station’ (Kema), now managed by the Institute of Tibetan Plateau Research, Chinese Academy of Science, southeast of Nagqu (Xizang Autonomous Region, 4450 m, 31°16’N / 92°06’E).

## DIVERSITY, DISTRIBUTION AND THE PALEO-ECOLOGICAL BACKGROUND

### Species diversity and distribution

The Tibetan highlands are a center of Cyperaceae diversity (Global *Carex* Group 2015), with more than 30 *Kobresia* species (Zhang and Noltie 2010). Recent studies have shown that the genus *Kobresia* should be included within *Carex* and that *K. pygmaea* should be called *Carex parvula* O. Yano (C.B. Clarke) (Global *Carex* Group 2015). However, given that these new proposals have not yet been implemented either in any of the locally relevant floras, nor in the ecological literature, the traditional nomenclature is retained here.

At altitudes between 3000 and 4000 m in the eastern highland forest-grassland ecotone, pastures of *Kobresia* species have developed: the height of the plants is between 10 and 20 cm, but the species vary, with *K. capillifolia* (Decne.) C.B. Clarke and *K. pusilla* N.A. Ivanova being dominant in the north; and *K. nepalensis* (Nees) Kük. in the south (Fig. 1). Sedge swamps along streams or in waterlogged areas over permafrost in the catchments of the Huang He, Yangtze, Mekong and Salween rivers are characterized by *K. tibetica* Maxim.; those along the Yarlung Zhangbo and Indus are composed of *K. schoenoides* Boeckeler – both species are between 15 and 60 cm high.

Two species, *K. pygmaea* and *K. yadongensis* Y.C. Yang reach hardly 4 cm in height. The latter has been described recently and its distribution is poorly known (Zhang and Noltie 2010, Miehe et al. 2011*a*). By far the most important species of the Tibetan rangelands is *K. pygmaea*. It was first collected in 1847 by Thomas Thomson in Rupshu (NW India), and is endemic to Tibet and the Inner Himalayas, ranging from the Deosai Plains of northern Pakistan to the Yulong Shan in northern Yunnan (Dickoré 1995, Zhang and Noltie 2010). The interspecific relationships between species as currently delimited remain unknown, because standard DNA for phylogenetic analysis (nuclear ITS and chloroplast DNA regions, including *mat*k, *trn*l-F and *trn*C-D) show low mutation rates and little divergence within *Kobresia* (J. Liu [*personal communication*]). The weak genetic differentiation suggests that these morphologically defined species may have evolved rapidly within the recent past. Further calibration, based on genomic data and fossils are needed to clarify evolutionary relationships among the *Kobresia* species as currently defined.

Vascular plant α-diversity of alpine *Kobresia* pastures (measured in plots of 100 m^2^) varies between 10 species in closed lawns with a *Kobresia pygmaea* cover of 98% (Miehe et al. 2008*b*, E. Seeber [*personal communication*]), and, in the eastern part of the plateau, more than 40 species in communities with mosaics of *Kobresia* patches and grasses, other sedges and perennial forbs growing as rosettes and cushions (Wang et al. 2017). Similar levels of richness were recorded in montane grazing lawns hosting a set of grazing weeds (Miehe et al. 2011*c*). The inter-annual variability of species richness is potentially high, depending on the variability of annual herbs in response to interannual changes in precipitation (E. Seeber [*personal communication*]).

In general, the Tibetan highlands, and specifically the eastern plateau, are poor in endemic plant genera and rich in endemic plant species (Wu et al. 1981). In contrast, and peculiar to the alpine *Kobresia* mosaics, is the high number of endemic monotypic genera of rosette plants, which colonize open soils around small mammal burrows (e.g., *Microcaryum pygmaeum* I.M. Johnst., *Przewalskia tangutica* Maxim., *Pomatosace filicula* Maxim.; Miehe and Miehe 2000).

The fauna also comprises several endemics, which are unevenly distributed among taxonomic and ecological groups. While most large predators are not endemic, a number of herbivorous mammals are, including wild yak, chiru and kiang, but also small mammals such as the plateau pika (Schaller 1998). In contrast, the beetle soil-fauna, which generally has a very high diversity in the Tibetan highlands (for example along wet gullies), is very poor and endemics are absent from the grazing lawns. Apparently the poor aeration of the turf, and the soil compaction due to trampling effects, are not suitable for the strictly edaphic ground beetle larvae (J. Schmidt [*personal communication*]). Important insects of the *Kobresia* pastures are Lepidoptera caterpillars of the genus *Gynaephora* (Erebidae, Lymantriinae; e.g., Yuan et al. 2015) which are known to cause severe damage (http://www.fao.org/ag/agp/agpc/doc/counprof/china/china2.htm, Yan et al. 1995, Xi et al. 2013, Zhang and Yuan 2013, Yuan et al. 2015). However, how regularly outbreaks occur, and how grazing intensity of *Kobresia* systems, or climatic effects, affect population dynamics of *Gynaephora* species are largely unknown. It appears that strongly grazed *Kobresia* ecosystems maintain only few phytophagous insects in terms of abundance and species richness – presumably because of the dominance of a single plant species, the high grazing intensity and the unfavorable environment. This coincides with the common finding that strong grazing by livestock decreases abundance and diversity of insects as caused by resource limitation, unfavorable microclimatic conditions and the lack of shelter due to reduced habitat heterogeneity (e.g., Littlewood 2008). Detailed studies across elevational gradients and across sites differing in grazing intensity are, however, needed to gain deeper insights as to how interactions with phytophagous insects affect the *Kobresia pygmaea* ecosystem in the long term.

### The paleo-ecological background

The development of biodiversity and endemism in relation to the highlands’ uplift and climate history is still under debate, hampering our understanding of the present diversity patterns. It is likely, however, that today’s evolutionary histories differ strongly between taxa and their traits such as climatic niches and dispersal ability. These two traits are especially relevant with respect Pleistocene climatic changes and likelihood of extinction. Table 1 reviews the available data on independent climate proxies for the Last Glacial Maximum (LGM; see Table 1). Most probably the world’s most elevated highlands were even more hostile for life during the cold periods of the Quaternary than today. The former idea of a complete ice cover across the highlands (Kuhle 2001) has, however, been soundly rejected on the basis of biotic and abiotic proxies. It was replaced by a concept of fragmented but locally extensive mountain glaciations, at least in the more humid eastern highlands (Shi 2006, Heyman et al. 2009).

**TABLE 1.** Estimates of temperature depression during the Last Glacial Maximum (maxDT) for the Tibetan highlands and the Central Himalaya. References are given in the Supplementary Material Table S1.

| Location | Method | maxDT [K] | Time [ka BP] | Reference |
| --- | --- | --- | --- | --- |
| Tibet Plateau | $\delta^{18}\text{O}$ value in ice core, ice wedges | 7 (annual) | 16-32 | Shi et al. 1997 |
| Guliya Ice Cap, West Kunlun | $\delta^{18}\text{O}$ value in ice core | 9 (annual) | 23 | Yao et al. 1997 |
| Dunde Ice Cap, Qilian Shan | $\delta^{18}\text{O}$ value in ice core | 6 (annual) | 30 | Thompson 1989 |
| Qarhan Salt Lake, Qaidam Basin | $\delta^{18}\text{O}$ and $\delta\text{D}$ of intercrystalline brine | 8 (annual) | 16–19.5 | Zhang et al. 1993 |
| Eastern Qilian Shan | Paleo-peat in frost heave | 7 (annual) | 31+/- 1.5 | Xu et al. 1984 |
| Gonghe Basin, Qinghai Province | Paleo-sand wedge | 7 (annual) | 17 +/- 0.25 | Xu et al. 1984 |
| Zoige Basin, East Tibet | Pollen analysis | 6 (annual) | 18 | Shen et al. 1996 |
| Zhabuye Lake, southwest Tibet | Pollen analysis | 6 (annual) | 18 | Xiao et al. 1996 |
| Hidden Lake & Ren Co, Southeast Tibet | Pollen analysis | 7–10 (January)/<br>0–1.5 (July) | 18–14 | Tang et al. 1999 |
| South and central Tibet | Atmospheric model | 3–5 (winter)/<br>2–4 (summer) | 21 | Liu et al. 2002 |
| High Asia | Estimate | <5 (mean summer) | LGM | Shi 2002 |
| Tibetan Plateau | Atmospheric model | 0.8–1.9 (annual) | 21 | Zheng et al. 2004 |
| High Asia | Atmospheric model | 6.3–6.4 (annual)/<br>5.6–6.1 (July) | 21 | Böhner and Lehmkuhl 2005 |
| Nyenquentangula Shan | Present elevational range of endemic beetles | 3–4 (summer temperature) | LGM | Schmidt et al. 2011 |
| Southern Tibet | Private juniper haplotypes | 3–4 (summer temperature) | LGM | Schmidt et al. 2011 |
| Changthang | Present elevational range of endemic plants | 3–4 (summer temperature) | LGM | Miehe et al. 2011 |
| Loess Plateau | Terrestrial mollusk | 3–5 (annual) | LGM | Wu et al. 2002 |
| Lake Rukche, Nepal | Forest/grassland biomarker | 3 (annual) | LGM | Glaser and Zech, 2005 |
| Kathmandu, Nepal | *Pinus*-Pollen | 3 (annual) | LGM | Paudyal and Ferguson 2004 |

Current paleo-scenarios for humidity and temperature still offer a wide range of possibilities, depending on the proxies on which they rely. The two most divergent paleo-scenarios – ecological stability or complete extinction and Holocene re-migration – can be assessed against the presence or absence of local endemics or populations with private haplotypes (Liu et al. 2014), because 10000 years since the LGM is too short a period for *in-situ* speciation.

In the case of the Tibetan highlands summer temperatures are particularly important. If these drop below minimum requirements for a given species, then local and endemic populations are lost – a phenomenon most important in closed-basin systems. The huge number of tiny, wingless, locally endemic ground beetles in the southern highlands is important in this respect, because it testifies to the persistence of alpine habitats throughout the Pleistocene and the absence of a glacial ‘tabula rasa’ (Schmidt 2011). The mean altitudinal lapse rate for summer temperatures across 85 climate stations (records between 1950 and 1980) in the highlands is 0.57 K per 100 m (Schmidt et al. 2011). Biogeographical data of the current distribution of these wingless ground beetles in the mountainous topography of southern Tibet point to a decline in LGM summer temperature of only 3–4 K as compared with present conditions (Schmidt et al. 2011), which is much lower than earlier estimates (Table 1). This is supported by estimates based on the presence of private haplotypes of juniper trees and endemic flowering plants in interior basins of the central Tibetan highlands (‘Changthang’; Miehe et al. 2011*b*, Schmidt et al. 2011), while records of terrestrial mollusks in loess deposited during the LGM on the Loess Plateau revealed annual temperatures 3–5 K lower than today (Wu et al. 2002). Biomarkers from the southern slope of the Himalaya (Lake Rukche, 3500 m; Glaser and Zech 2005) indicate a downward shift of the upper tree line of 500 m, similar to shifts of distribution boundaries of two *Pinus* species in the Kathmandu basin of Nepal (Paudayal and Ferguson 2004). These also lead to an estimated LGM temperature drop of only 3–4 K (Miehe et al. 2015).

The development of humidity scenarios for the LGM may also benefit from applying ecological indicator values of endemic plants or paleo-pollen records. For example, the persistence of juniper trees in Tibet (Opgenoorth et al. 2010) throughout the LGM implies that rainfall was never lower than 200 mm/yr, because the present drought line of junipers correlates with 200–250 mm/yr. Grass pollen records in the north-eastern highlands (Lake Luanhaizi; Herzschuh et al. 2006) and in southern Tibet (Nangla Lu; Miehe et al. [*personal communication*]) reveal the presence of steppe vegetation and thus an annual rainfall of more than 100 to 150 mm. However, the true values may be somewhat lower as colder climates would result in less evapotranspiration. The LGM– loess records in Tibet (Kaiser et al. 2009*a*) also testify to the presence of grass cover as a precondition for loess accumulation. Thus, both temperature and humidity during the LGM allowed the persistence of plants, herbivores and hunters in the Tibetan highlands.

## LIFE HISTORY TRAITS AND REPRODUCTION OF *KOBRESIA PYGMAEA*

*Kobresia pygmaea* is one of the smallest alpine sedges, yet dominates the largest alpine pastoral ecosystem, stretching over an altitudinal range of 3000 m. Its lawns have an estimated Leaf Area Index (LAI) of only ∼1 (Hu et al. 2009) and a roughness length of only about 3 mm. They represent a vegetation cover with a small transpiring surface and low aerodynamic resistance to the atmosphere (Babel et al. 2014). Due to high solar radiation input and night-time long-wave radiation, *Kobresia pygmaea* has to cope with steep soil-to-air temperature gradients and high leaf over-temperatures on sunny days. However, the species shows low sensitivity to low soil temperatures, as leaf gas exchange was found to be negatively affected only by soil temperatures below freezing point, i.e., when soil-water availability approaches zero (Coners et al. 2016).

In fully sun-exposed sites, *Kobresia pygmaea* grows mostly in lawns of 2 cm in height, yet lawns of only 1 cm or up to 4 cm can occur locally, regardless of being grazed or ungrazed, and whether growing in water-surplus sites such as swamps or on steep, dry slopes. The phenotypic similarity across the whole range of environmental conditions in the Tibetan highlands shows the species’ adaptation to the cold and seasonally dry environment, and is probably one of the factors explaining its wide distribution in the highlands. Grazing exclusion experiments, however, have revealed that *K. pygmaea* can attain 20 cm when overgrown and shaded by taller grasses or under tree crowns (S. Miehe and Y. Wang [*personal communication*]). This plasticity may explain the extraordinary wide range of distribution and is possibly an effect of polyploidy. The same effect was observed in a plant growth chamber with climatic parameters closely simulating the conditions of a typical summer day at the alpine core range southeast of Nagqu (Kema site, 4450 m; H. Coners [*personal communication*]). Although the lamps used were chosen for their spectral similarity to sunlight, after some weeks *Kobresia pygmaea* plants reached up to 20 cm in height. Artificial lighting cannot reach the peak intensities in radiation pertaining on the plateau (especially in the UV-B part of the spectrum), which are among the highest occurring on earth (Beckmann et al. 2014). Whether permanently high radiation loads represent an inhibitor with a photo-toxic effect that reduces the growth of *Kobresia*, remains unclear; the same is true for effects of wind and desiccation. No doubt, the dwarf habit could also have evolved as an effective strategy to avoid plant biomass loss and damage by grazing.

A microsatellite-based survey of alpine *Kobresia pygmaea* populations revealed the presence of large (>2 m^2^) clones, and an overall high genetic diversity (with more than 10 genets/m^2^). This probably results from consecutive events of sexual recruitment under favorable conditions, coupled with extensive periods of vegetative persistence (Seeber et al. 2016). Thus, in contrast to many alpine species that largely abandon sexual reproduction (Steinger et al. 1996, Bauert et al. 1998), *K. pygmaea* benefits from a mixed reproduction strategy with clonal growth ensuring long-term persistence and competitiveness, while intermittent reseeding facilitates colonization and genetic recombination. A diaspore bank supports short-term persistence. According to microsatellite studies, populations are hardly separated between montane and alpine altitudes, indicating high gene flow within the species’ distribution range (Seeber 2015). The genetic diversity in the species is also promoted by polyploidy; *K. pygmaea* mostly has 2*n* = 4*x* = 64 chromosomes and is tetraploid. Ploidy of congeners range from di- to hexa- or even heptaploidy (Seeber et al. 2014). An assessment of ploidy levels along an elevational gradient in Qinghai also revealed one diploid, one triploid, two octoploid and one dodecaploid individual, indicating ongoing chromosomal genetic evolution in *K. pygmaea* (E. Seeber [*personal communication*]).

*Kobresia pygmaea* plants are mostly monoecious, with androgynous spikes (upper flowers male, lower female; Zhang and Noltie 2010), yet dioecious forms do occur (own observations at 4200 m, 33°12’N/97°25’E, plants with entirely male spikes). High numbers of inflorescences are produced (∼100 to 5000 inflorescences/m^2^; Seeber et al. 2016), depending on the weather conditions in the respective year but not on the grazing regimes. Diaspores are highly viable and adapted to (endo)-zoochorous dispersal. Germination rates differ greatly between seeds from alpine and montane environments (e.g., Li et al. 1996, Deng et al. 2002, Miao et al. 2008, Huang et al. 2009, Seeber et al. 2016) but are generally low, both in laboratory experiments as well as *in situ*. Huang et al. (2009) reported 13% germination of untreated diaspores, while most studies obtained no seedlings at all (Li et al. 1996, Miao et al. 2008, Seeber et al. 2016). The water-impermeable pericarp hinders germination, and its removal by chemical or mechanical interventions increases germination rates. Under natural conditions, microbial activity or digestion by herbivores may have the same effect and, consequently, increase germination. Nonetheless, the variability in germination was high between different studies, which may reflect population responses to different abiotic conditions, such as nutrient availability or elevation (Amen 1966, Seeber 2015).

## WATER BUDGET AND HYDROLOGICAL FLUXES OF ECOSYSTEMS

*Kobresia* pastures exist over a wide range of precipitation from 300 to 1000 mm/yr (Miehe et al. 2008*b*), which falls nearly exclusively during summer. Apart from nutrient limitation (see below), the growth of *Kobresia pygmaea* is co-limited by low summer rainfall, at least under the conditions of the ecosystem’s alpine core range (Kema; Coners et al. 2016) with 430 mm/yr and mean maximum temperature of the warmest month of 9.0 °C (1971 – 2000; Babel et al. 2014). Onset of the growing season is controlled by rainfall amount in early summer and under low-temperature control in autumn (September and October, depending on latitude and altitude). The leaf growth of *K. pygmaea* is not temperature-driven (unlike that of *Androsace tapete* Maxim., a cushion plant of the alpine steppe), but depends strictly on water availability (Li et al. 2016). Greening of the *Kobresia* pastures after the onset of the summer rains is well known; this usually occurs between mid-May and mid-June (Fig. 2). However, onset of the summer monsoon can be delayed by up to six weeks, sometimes starting as late as early August, with critical effects for livestock (Miehe and Miehe 2000).

**FIG. 2.**
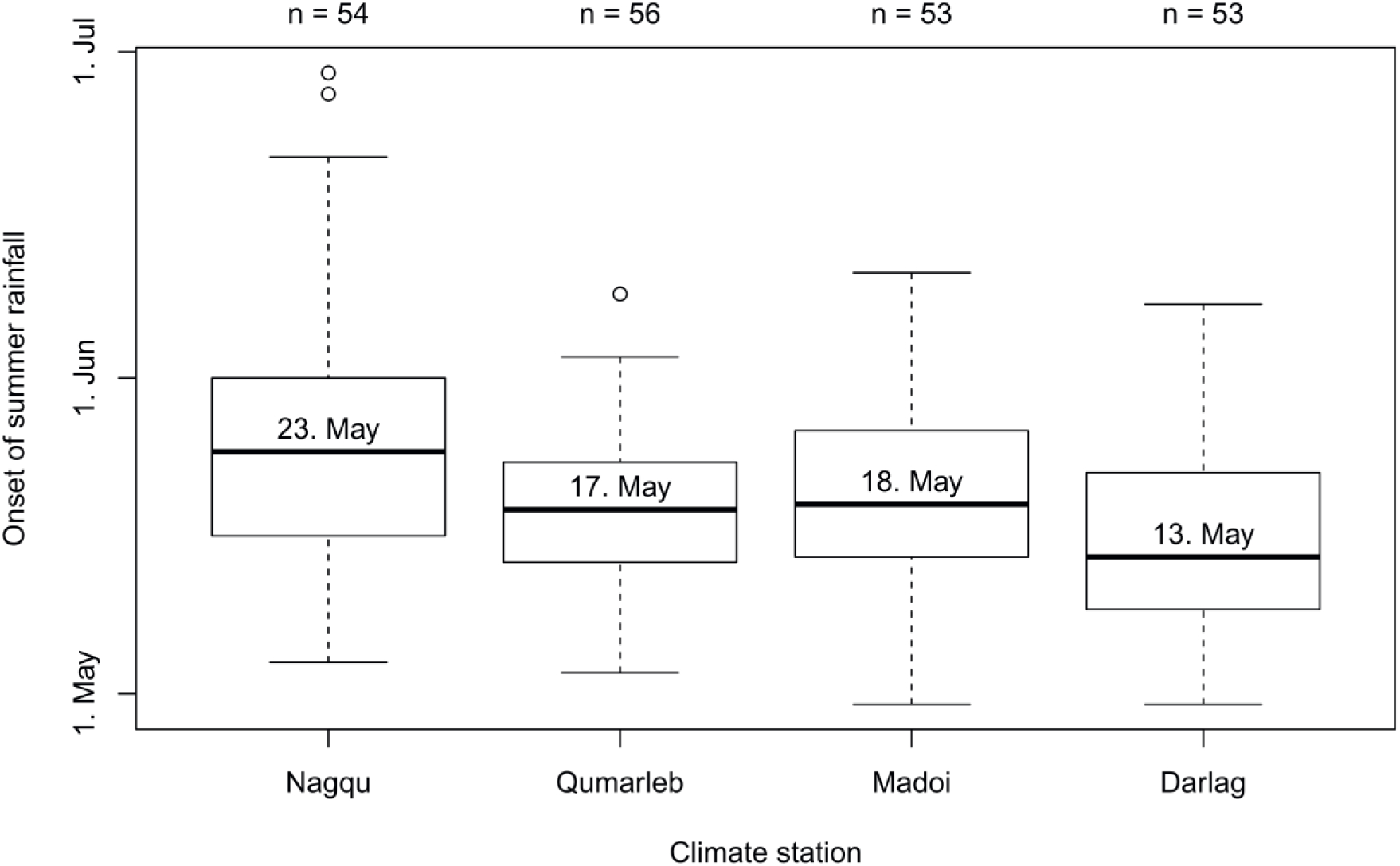
Date of the onset of the summer precipitation observed at selected climate stations. In the boxplots, lines and boxes correspond to extreme values, the first quartile and the third quartile. The written date is the median (black solid line in the boxplots) of the onset for the respective station (n is the number of years with sufficient observations). For the calculation of the onset, the first derivation of a 6^th^ order polynomial fitted between daily precipitation sums and the day of the year is derived for each year. The date of the onset is defined as the date when the maximum of the first derivation occurred. Precipitation data is from the Global Historical Climatology Network (GHCN-d; Menne et al. 2012).

Despite its low leaf area index, alpine *Kobresia* pastures can reach high transpiration rates (up to 5 mm/day) in moist summer periods even at elevations >4000 m: a consequence of specific microclimatic conditions on the plateau which enhance evaporation (Coners et al. 2016). While dry periods, due to delayed onset of the summer rains, visibly hamper *K. pygmaea* growth (browning of the pastures), a constant daily irrigation with 2.5 mm/day or even 5 mm /day did not increase the total above-ground biomass production after 40–70 days (corresponding to 100–350 mm of added water; Coners et al. [*personal communication*]). This matches observations of constant biomass production despite large variation in precipitation (Climatic station of Nagqu 2011: 574 mm, 2012: 437 mm, 1971–2000: 430 mm; Seeber 2015, http://www.geodata.us/weather/index.html, accessed January 2015). Lysimeter experiments show that the evapotranspiration closely depends on precipitation. Most of the atmospheric water input, except for heavy rainfall, is lost immediately by evapotranspiration (Coners et al. 2016).

Due to the highland’s relevance for global circulation changes, the surface properties have impacts on large, and possibly even global scale. In terms of landscape-level moisture cycles, pasture degradation leads to a shift from transpiration to evaporation because of reduced biomass, although the evapotranspiration changes are not significant over a longer period (Babel et al. 2014). Modelling indicates an earlier onset of convection and cloud generation, probably triggered by a shift in evapotranspiration timing when dominated by evaporation. Consequently, precipitation starts earlier and clouds decrease the incoming solar radiation. Thus changes in surface properties due to pasture degradation have a significant influence on larger scales with respect to the starting time of convection and cloud- and precipitation-generation: convection above a degraded surface occurs before rather than after noon. Due to the dominant direct solar radiation on the Tibetan highlands the early-generated cloud cover reduces the energy input and therefore the surface temperature (Babel et al. 2014). This can have negative effects on the intensity of plant photosynthesis, and thereby reduces the ecosystem’s ability to recover from degradation. The changes in the water cycle are furthermore influenced by global warming and an extended growing season (Fig. 3; Che et al. 2014, Shen et al. 2014, Yang et al. 2014). In many years however, effects are overtaken by the water availability of the delayed onset of the summer rains.

**FIG. 3.**
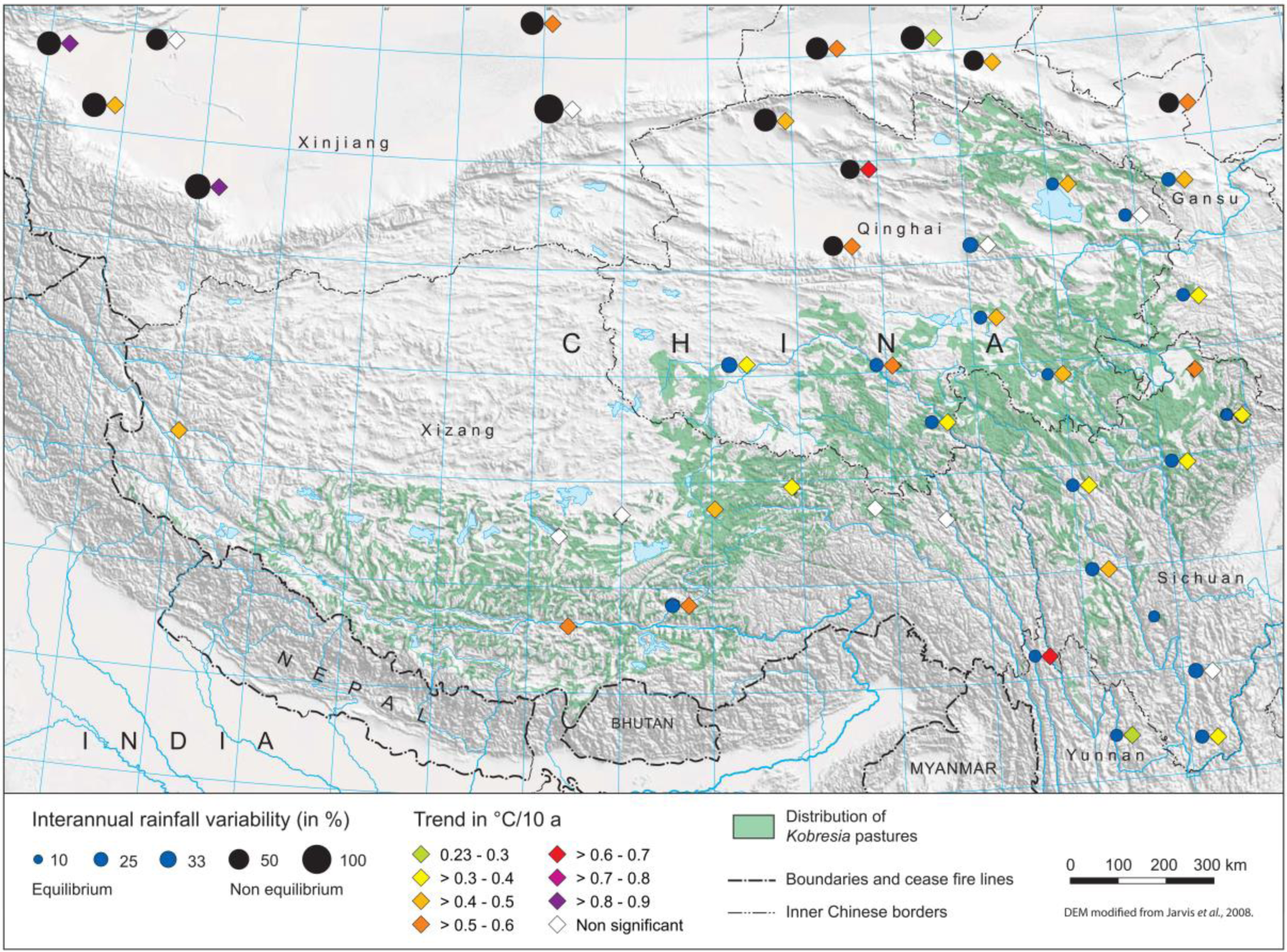
Inter-annual variability of summer rainfall sums defined as the standard deviation of summer rainfall sums divided by the mean summer rainfall sum across the Tibetan Plateau (June, July, August) between 1986 and 2015, and trends in summer mean temperatures (June, July, August) between 1985 and 2015 based on daily stations’ observations from the Global Historical Climatology Network (GHCN-d; Menne et al. 2012). The trend is calculated from linear models and significance is derived using Mann-Kendall correlation techniques.

The status of the *Kobresia* cover and its root mat is nowhere of greater importance than in the permafrost areas of the Salween, Yangtze and Huang He headwaters, covering an area of approximately 180000 km^2^. The turf’s insulating effect buffers the melting of the permafrost; soil temperatures under a patchy vegetation cover of 30% were found to be 2.5 K higher than under a closed mat of 93% (Wang et al. 2008). The recent (1980–2005) increase in surface soil temperatures in the Huang He headwaters of 0.6 K per decade has led to a drastic increase of the depth of the thawing layer (Xue et al. 2009), and to a deterioration of *Kobresia tibetica* swamps. In summary, overgrazing induces degradation of root mats and leads to changes in water cycle and balance at both local and regional levels; this may decrease the recovery of damaged *Kobresia* pastures.

## THE CARBON CYCLE

*Kobresia* turf is a key component of the C stocks and cycle in these pastures. The root : shoot ratio ranges from 20 (Li et al. 2008, Unteregelsbacher et al. 2012, Schleuss et al. 2015) up to 90 (Ingrisch et al. 2015), depending on season, grazing intensity, and degradation stage. Soil organic carbon (SOC) storage within the root mat reaches up to 10 kg C/m^2^ (Li et al. 2008, Unteregelsbacher et al. 2012, Schleuss et al. 2015), making up roughly 50% of the overall C stocks. Representing a highly significant C stock, the turf is also a highly active component of the C cycle, which receives the largest fraction of the photo assimilated C. For alpine *Kobresia* pastures at 4450 m, Ingrisch et al. (2015) showed that a large fraction of the assimilates is used for the build-up of new fine roots with a high turnover rate. The measured fluxes into below-ground pools, mainly associated with the root and released later as soil CO_2_ efflux, were roughly twice as high as reported for pastures of *K. humilis* (C.A.Mey. ex Trautv.) Serg. (Wu et al. 2010) and montane *K. pygmaea* pastures at lower elevation (3440 m; Hafner et al. 2012). This emphasizes the importance of below-ground C allocation and cycling in alpine *K. pygmaea* pastures.

Net ecosystem exchange (NEE) measurements, derived by the eddy-covariance method, identified the alpine pastures at Kema as a weak C sink during the summer of 2010 with an ecosystem assimilation of 1.3 g C m^-2^ d^-1^ (Ingrisch et al. 2015), which is roughly 50% smaller than recorded in pastures at lower altitudes (e.g., Kato et al. 2004, 2006, Zhao et al. 2006, Hirota et al. 2009), but agrees well with the results of a study by Fu et al. (2009) at a similar altitude. By combining the NEE measurements with ^13^C - pulse labelling, Ingrisch et al. (2015) were able to estimate absolute C fluxes into the different ecosystem compartments during the main growing season. With a magnitude of 1 g C m^-2^ d^-1^, the flux into below-ground pools was twice as high as the CO_2_ efflux from soil, and 10 times larger than the C flux into the above-ground biomass.

The key role of grazing for C sequestration and C cycling was demonstrated in two grazing exclusion experiments combined with ^13^CO_2_ pulse labelling studies. The effect of grazing on the C cycle, specifically on differences in below-ground C stocks and C allocation, was shown for:

1. a montane *Kobresia* winter pasture of yaks, with moderate grazing regime, compared to a 7-year-old grazing exclosure plot, both at 3440 m (Xinghai; Hafner et al. 2012), and
2. an alpine *Kobresia* pasture compared to a 1-year-old grazing exclosure, both at 4450 m (Kema; Ingrisch et al. 2015).

Short-term grazing exclusion in the alpine pasture affected only the phytomass of above-ground shoots, while neither C stocks nor assimilate allocation were altered (Ingrisch et al. 2015). In this system, roots and soil were of equal importance to C storage. By contrast, 7 years of grazing exclosure revealed that grazing is a major driver for below-ground C allocation and C sequestration in soils of montane *Kobresia* pastures (Hafner et al. 2012). Under a grazing regime, a higher fraction of assimilated C was allocated to below-ground pools; moreover, a larger amount was incorporated into roots and SOC. Fencing in contrast, led to a significant reduction of C sequestration in the soil and fostered turf degradation, emphasizing the key role of grazing for the biogeochemistry of these ecosystems.

Based on the long-term grazing exclosure experiments in montane pastures, we conclude that the larger below-ground C allocation of plants, the larger amount of recently assimilated C remaining in the soil, and the lower soil organic-matter derived CO_2_ efflux create a positive effect of moderate grazing on soil C input and C sequestration in the whole ecosystem. Due to the large size of the below-ground C stocks and the low productivity of the ecosystem, changes in the C stocks after cessation of grazing can be expected to take at least several years to become apparent. However, the roots in the turf mat are a highly dynamic component of the C cycle, which might have implications for the interannual variability of the C budget even on the landscape scale. The C cycle appears to be largely driven by grazing, supporting the hypothesis of the pastoral origin of the *Kobresia* ecosystem.

At Kema research station, synchronous measurements with micro-lysimeters, gas exchange chambers, ^13^C labelling, and eddy-covariance towers were combined with land-surface and atmospheric models, adapted to the highland conditions. This allowed analysis of how surface properties, notably the disintegration of the *Kobresia* sward (i.e., degradation stages), affect the water and C cycle of pastures on the landscape scale within this core region. The removal of the *Kobresia* turf fundamentally alters the C cycling in this alpine ecosystem and its capacity of acting as a C sink (Babel et al. 2014).

## SOILS, PRODUCTIVITY, AND PLANT NUTRITION

*Kobresia* lawns typically produce a felty root mat (Afe, ‘rhizomull’; Kaiser et al. 2008) up to 30 cm thick, which is situated on top of the predominant soils of Tibet’s pasture ecosystems – Leptosols, Kastanozems, Regosols, Cambisols and Calcisols (Fig. 4; Kaiser 2008, Kaiser et al. 2008, Baumann et al. 2009). The root mat consists of mineral particles, humified organic matter and large amounts of dead and living roots as well as rhizomes (Schleuss et al. 2015) The mat has typically formed in a loess layer of Holocene age (Lehmkuhl et al. 2000). The question still remains as to whether the *Kobresia* lawns and their root mat sealed a pre-existing Ah-horizon of a tall grassland, or if the lawns have grown up with the loess that they have accumulated (Fig. 4; Kaiser et al. 2008).

**FIG. 4.**
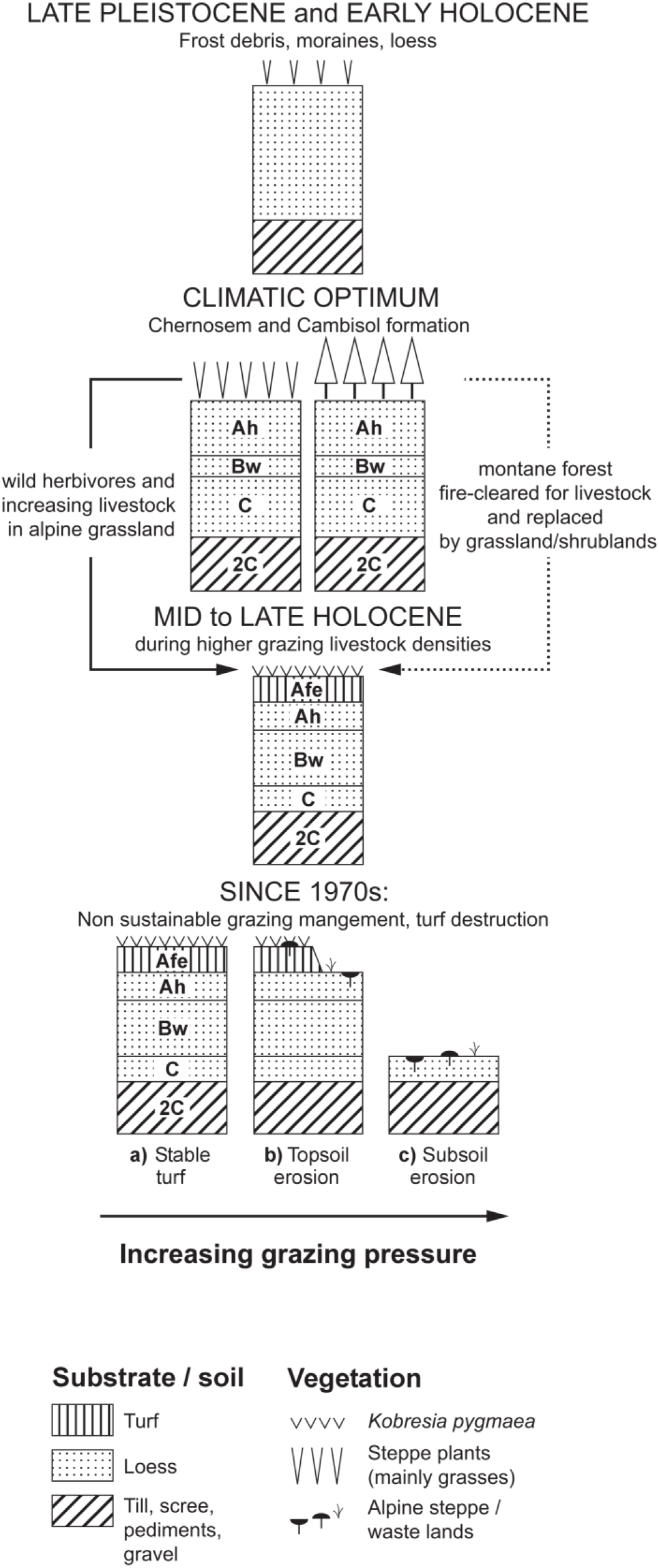
Hypothetical dynamics of soils in alpine grasslands of the southeastern Tibetan highlands. After Kaiser et al. 2008, Miehe et al. 2011*c*.

At the core alpine Kema study site, the root-mat is characterized by low bulk densities, moderate pH values, high SOC contents and a high effective cation exchange capacity (CEC), of which calcium (Ca^2+^) is the most abundant base cation (up to 80% of the effective CEC; Fig. 5). The root mat stores about 1 kg N/m^2^ and 0.15 kg P/m^2^ (0–30 cm; Fig. 5), but most nutrients are stored in soil organic matter (SOM) and dead roots and are not directly plant-available. Prevailing low temperatures and moisture hamper the mineralization of root residues and slow down nutrient release to plant-available forms (Hobbie et al. 2002, Luo et al. 2004, Vitousek et al. 2010). From this perspective, the relatively close soil C/N and C/P ratios (Fig. 5) do not necessarily indicate an adequate N and P supply. Further, P can be precipitated in the form of calcium-phosphates due to the high abundance of exchangeable Ca.

**FIG. 5.**
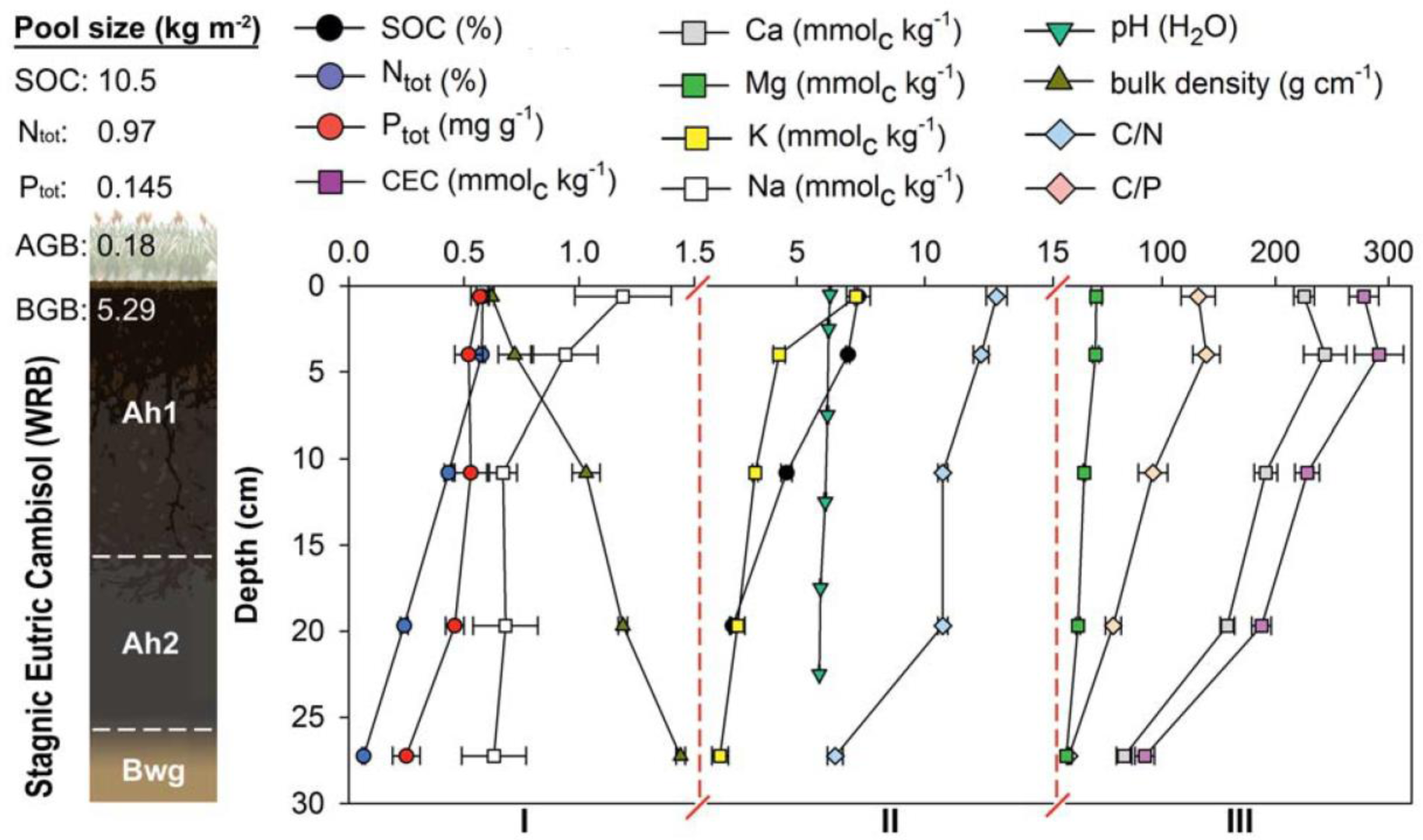
Basic characteristics of root mats at the Kema research sites in the *Kobresia pygmaea* core area. Pool sizes include below-ground-biomass (BGB), stocks of soil organic carbon (SOC), total nitrogen (N_tot_) and total phosphorus (P_tot_) down to 30 cm, and aboveground biomass (AGB). Note that the soil physical and chemical properties are plotted on three different scales (I: 0–1.5, II: 1.5–16, III 16–300). Data are represented as means (*n* = 4) and error bars indicate standard errors.

Indeed multiple limitations of N and P constrain pasture productivity, which is shown by increased *Kobresia* growth following single and combined applications of N and P fertilizers. Even though single applications of either N or P favor above-ground biomass on most sites throughout the *Kobresia* ecosystem, the productivity strongly increases after combined N and P application (Fig. 6). This finding was also supported by three years of single and combined application of potassium (K), N and P in alpine *Kobresia* pastures at the Kema station. Nitrogen fertilization increased above-ground productivity about 1.2–1.6 times, while NP addition resulted in 1.5–2.4 times higher values, whereas no effect was found for the below-ground biomass (Seeber 2015). Furthermore, fertilization increased the tissue content of N, P and K in *K. pygmaea* and in accompanying herbs. Overall, fertilization experiments clearly indicate that co-limitations of N and P prevail in the *Kobresia* ecosystem. We conclude that *Kobresia pygmaea* has developed a dense root network to cope with nutrient limitations enabling productivity and competitive ability. The high below-ground biomass on the one hand ensures an efficient uptake of nutrients (shown by ^15^N; Schleuss et al. 2015) at depths and times when nutrients are released via decomposition of SOM and dead roots; and on the other hand makes *K. pygmaea* highly competitive for mineral N acquisition in comparison with other plant species (Song et al. 2007) and microorganisms (Xu et al. 2011, Kuzyakov and Xu 2013). Further, the enormous root biomass stores nutrients below-ground, protecting them from removal via grazing, which ensures fast regrowth following grazing events to cover the high below-ground C costs.

**FIG. 6.**
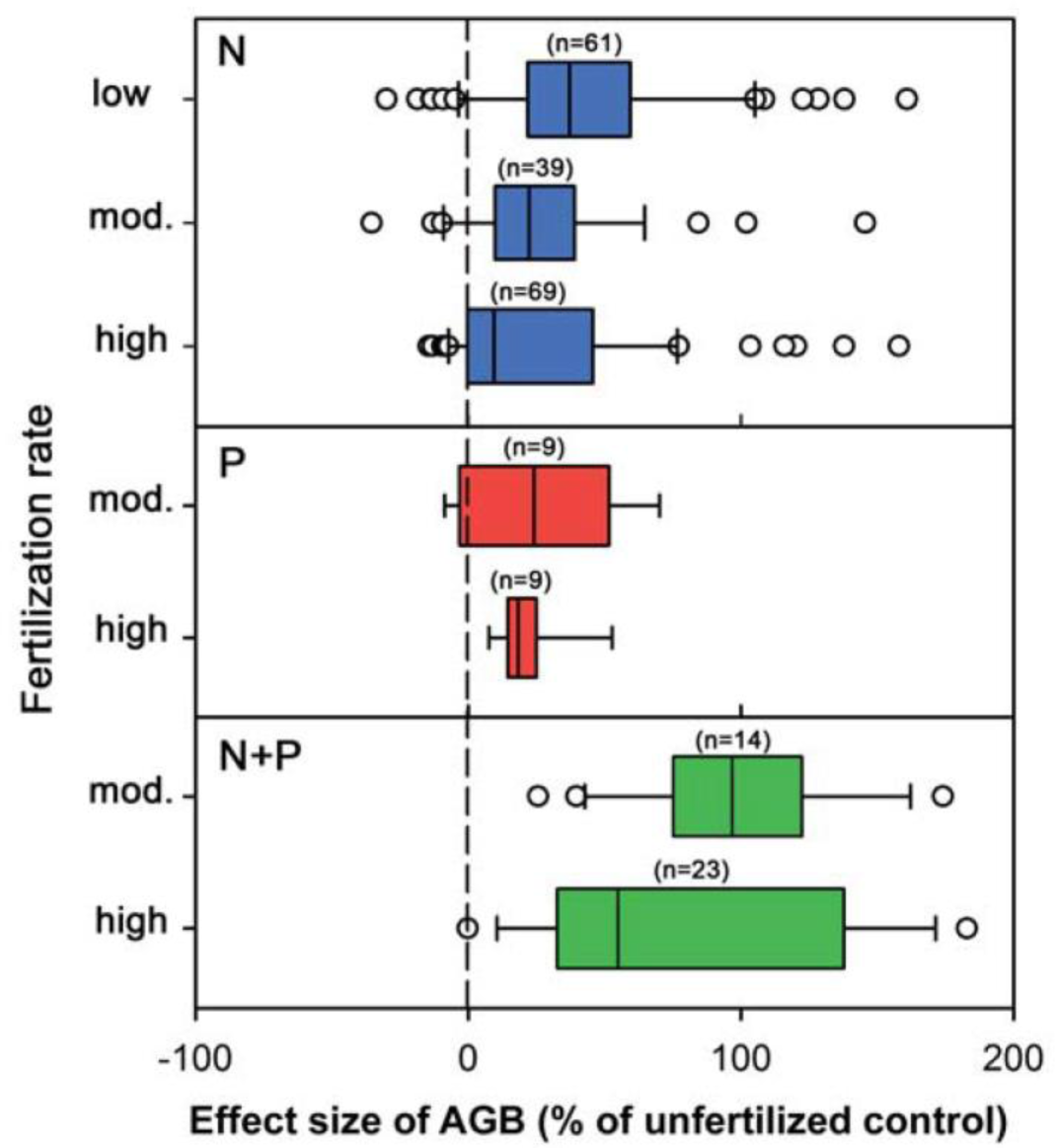
Effects of single and combined fertilization with N and P at varying rates on aboveground biomass (AGB) extracted from 35 studies from all over the *Kobresia* ecosystem. Shown are Whisker-Box-Plots with outliers (white circles) and median (black line in the box) for low, moderate and high application rates (for N: low = 0–25 kg/ha*yr, moderate = 25–50 kg/ha*yr, high >50 kg/ha*yr; for P: moderate = 0–50 kg/ha*yr, high >50 kg/ha*yr; for N+P: moderate = 0–50 + 0–50 kg/ha*yr, high >50+50 kg/ha*yr). The dashed line indicates no effects, with negative effects on the left and positive effects following fertilization on the right.

That considerable amounts of resources were allocated and stored below-ground was confirmed by labelling studies showing that about 45% of ^13^C (after 48 days) and 50% of ^15^N (after 45 days) were transferred into root biomass (0–15 cm; Ingrisch et al. 2015, Schleuss et al. 2015).

In alpine *Kobresia* pastures around Kema the annual above-ground biomass productivity is low (∼132 g/m^2^, S. Träger and E. Seeber [*personal communication*]) compared to below-ground plant biomass (0–15 cm horizon about 5–6 kg/m^2^; Seeber 2015). This corresponds well with other sedge-dominated montane and alpine Tibetan grasslands (134 g/m^2^; Ma et al. 2010) but is more than double that of the above-ground productivity of alpine steppes in the arid north-western highlands (56 g/m^2^).

Comparing montane (Xinghai) with alpine pastures (Kema) also revealed clear differences in above-ground productivity with altitude (montane: 185 g/m^2^, alpine: 132 g/m^2^; S. Träger and E. Seeber [*unpublished manuscript*]). Presumably the higher temperatures in Xinghai (Babel et al. 2014), together with a longer vegetation period and higher nutrient availability (N, P) due to enhanced mineralization rates in warmer sites, resulted in increased biomass production in montane as compared with alpine sites.

The above results refer to the scale of soil profiles and individual plots, and neither consider the redistribution of nutrients within the very heterogeneous landscape, nor account for the reallocation of nutrients by intensive grazing and animal movement. On a landscape scale the striking contrast between the widespread, chlorotic, yellowish *Kobresia* mats and the localized, bright-green cattle resting places around settlements points to the livestock-mediated nutrient translocation that has been described in Chinese and Mongolian steppes (Stumpp et al. 2005, Holst et al. 2007, Wesche and Ronnenberg 2010). Yak dung is the exclusive and indispensable fuel for highland nomads (Rhode et al. 2007), and dung collection thus has resulted in nutrient export since the onset of pastoralism, i.e., since the mid-Holocene up to 8000 year BP (Miehe et al. 2014, see below). Increased stocking rates and reduced migration distances since the late 1970s (Zhou et al. 2005) have certainly aggravated naturally existing gradients in nutrient availability with very high concentrations around villages.

## PASTURE HEALTH AND DEGRADATION

The term ‘degradation’ can refer not only to widespread negative effects of rangeland management, but also to natural processes of ecosystem disturbance that are often poorly understood. On the Tibetan highlands, degradation is by no means equally distributed; it is more severe (1) in the vicinity of settlements, (2) on the lower slopes of southern exposures and (3) in the ecotone areas between steppes and alpine meadows with moderate rainfall (Miehe and Miehe 2000, Wang et al. 2017).

The *Kobresia* ecosystem is an equilibrium grazing system with a coefficient of variance in interannual rainfall variability of well below 30% (Fig. 3; Ellis and Swift 1988, Ellis 1995, von Wehrden et al. 2012). In contrast to more variable (semi-)arid, non-equilibrium systems, grazing impact is not regularly set back by largescale loss of livestock caused by shortage of rainfall and thus forage. With their relatively stable forage resources, equilibrium systems may degrade if livestock numbers increase until the carrying capacity is exceeded. The impact of severe snowstorms that occur irregularly and regionally on the plateau and also lead to losses of livestock (Yeh et al. 2014), has not, however, yet been analyzed with respect to livestock dynamics. Snowstorms introduce another form of climate variability and may prevent livestock from increasing beyond the carrying capacity. The question, therefore, is whether the alpine grazing lawns are as vulnerable as other equilibrium pasture systems. Indeed, the specific traits of the prevailing species described above suggest that degradation threat may be limited.

Estimates of grazing-induced degradation vary: the most frequently quoted value for the Tibetan highlands is that 30% of the grasslands are degraded (Harris 2010, Wang and Wesche 2016). The loss in ecosystem services caused by the C emission and N export associated with pasture degradation has been calculated to amount to $8033/ha and $13315/ha respectively (Wen et al. 2013) in the Dawu area. In the Ruoergai Plateau, also known as Zoige Plateau, the ecosystem services value expressed as a multiple of the gross domestic product (GDP) has decreased by about 84% between 1990 and 2005 (Li et al. 2010). Inconsistent definitions, unclear baselines, varying standards and indicator systems, as well as the merging of different spatial scales, result in uncertainties in these calculations. Desertification is often not differentiated from degradation (Wang et al. 2008, Cui and Graf 2009) and, in general, climate change and human impact are rarely separated (Chen et al. 2014, Zhou et al. 2014, Fassnacht et al. 2015, cf. He et al. 2015).

Remote sensing may offer an option for large-scale assessments of degradation, yet suffers from drawbacks. Degradation itself cannot be detected directly, and remote sensing requires extensive ground-truthing to calibrate and validate the results. Thus remote sensing can detect changes in degradation only over time, as it has been demonstrated by Lehnert et al. (2016) based on an area-wide plant cover dataset (Lehnert et al. 2015). The results show that degradation since the year 2000 is proceeding only in the less productive *Kobresia* pastures in the western part of the highlands, where it is largely driven by slight decreases in precipitation, in combination with rising temperatures. In contradiction to the widely assumed high importance of human influence on the degradation process, stocking numbers were not strongly correlated with larger-scale plant cover changes (Lehnert et al. 2016).

Difficulties in properly selecting and interpreting spectral data may also render large-scale remote-sensing-based assessments and models questionable (Yang et al. 2005). Fine-scale changes in vegetation and soils represent appropriate indicators for local, site-based assessments, but they are not easily detected by spectral data, and are not representative for the entire highland region. Thus pasture degradation is clearly a phenomenon with diverging regional gradients, depending on climate, soils and the regionally different impacts of rangeland management change.

Traditional nomadic systems cope with environmental heterogeneity and variability in resource availability by conducting seasonal migratory and other movements. Since the 1960s, government interventions have changed rangeland policies and led to an increase of sheep and goats by 100% in the early 1980s, causing severe damage to rangelands regionally (Zhou et al. 2005). State regulations nowadays focus on destocking, sedentarization, privatization and the fencing of pastures, thereby reducing the mobility and flexibility of the herders, with potentially severe consequences for the development of pastures and an increased threat of degradation (Qiu 2016). To maintain the ecosystem’s services the development and implementation of scientifically proven, and regionally adapted, modern rangeland management systems are necessary.

A number of studies addressed the issue of degradation from a soil perspective. Under optimum conditions, *Kobresia pygmaea* builds almost closed, mono-specific, golf-course like lawns in high altitudes above 4600–4900 m (Fig. 7A and B). More common, however, are patterns showing degradation phenomena of uncertain origin and dynamics. The most widespread are (1) polygonal crack patterns (Fig. 7C) and the drifting downwards of polygonal sods (Fig. 7D), (2) sods resting like stepping-stones on a deflation-pavement, otherwise covered with alpine steppe plants (Fig. 7E, F), and (3) patchwise dieback of the lawn in front of the burrows of soil-dwelling small mammals (e.g., *Ochotona curzoniae* Hodgson, pika; Fig. 7H, J). All three patterns can be observed across the entire range of the ecosystem, but have been most well documented in an ecotone stretching 200 km in width over 2000 km across the whole highlands between the Qilian Shan in the north and the Himalayas in the south (Miehe (1) et al. 2011*c*). Polygonal cracks are found all over the range of *Kobresia* lawns, but they occur only in root mats of more than 5 cm in thickness. Interviews with herders confirm that low winter temperatures cause the cracking, but there also is evidence of desiccation effects (Miehe and Miehe 2000, Schleuss et al. [*personal communication*]). Both extreme events cause changes in the volume of the sods. As soon as the root mat has reached a certain thickness with a large portion of dead roots, tensions resulting from the volume changes lead to the formation of polygonal cracks. Overgrazing and trampling may play an additional role in weakening the stability of the root mat. On the prevailing steep inclination, polygons are progressively separated while drifting downhill with gravity, frequently above a wet and frozen soil layer (Fig. 7D). The widening of the cracks is accompanied by high SOC losses (∼5 kg C/m^2^). Moreover, SOC loss is aggravated by decreasing root C input following root decay and SOC mineralization indicated by decreasing SOC contents with intensified degradation. A negative δ^13^C shift of SOC caused by the relative enrichment of ^13^C-depleted lignin confirmed this mineralization-derived SOC loss (∼2.5 kg C/m^2^). Overall up, to 70% of the SOC stock (0 cm to 30 cm) was lost in comparison with intact swards of alpine *Kobresia* pastures in the Kema region (Schleuss et al. [*personal communication*]). Here, a degradation survey revealed that about 20% of the surface area has lost its *Kobresia* turf with bare soil patches remaining (Babel et al. 2014). Assuming that the whole *Kobresia* ecosystem has suffered from this type of degradation to a similar extent, and that the soil conditions in Kema (SOC stock ca. 10 kg m^-2^)) are representative across the highlands, this would imply a total SOC loss of 0.6 Pg C for the whole ecosystem with its 450000 km^2^. Consequently a high amount of C is released back to the atmosphere as CO_2_, or is deposited in depressions and rivers – leading to a decline of water quality at both landscape and regional levels.

**FIG. 7.**
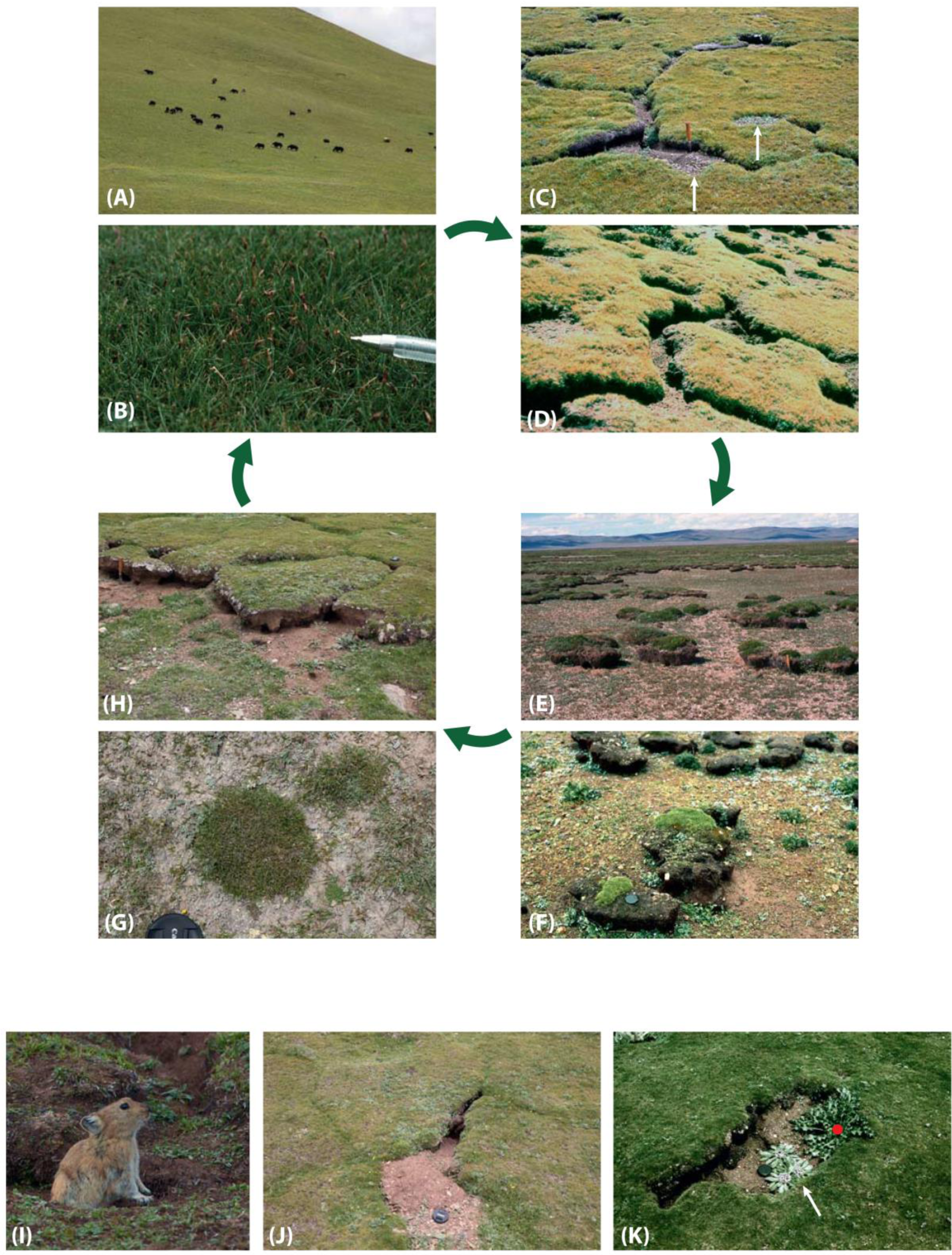
Autocyclic model of turf degradation in *Kobresia pygmaea* pastures. (AB) Closed grazing lawns are the best yak pastures. (C) Polygonal separation of the felty root mat, and D) downslope drift of the sods. (EF) The former turf cover is destroyed into stepping-stone like relics. The turf cliffs of 25 cm in height are corroded by needle-ice, wind and undercut by pika excavations. The surrounding open soil and gravel carry alpine steppe species. (GH) Recolonization of pancake-like *Kobresia pygmaea* mats in the open soil in front of the turf-cliff. (IJK) Pikas increase habitat diversity mainly through their digging; the excavated soil covers the lawns and root mats (J) A patchwise dieback of lawns and erosion of the felty root mat follows (*cf*. arrows in Fig. C). (K) The open soil is colonized by endemic annuals *Microula tibetica* (arrow) and *Przewalskia tangutica* (red dot). Photos G. Miehe 1994–2015.

The widening space between the crack margins is frequently, but not exclusively, used by pikas to dig their burrows, and they also undermine the ‘cliffs’ for deposition of their faeces. Excavated soil covers the lawns in front of their burrows (white arrows in Figs. 7C, J), which leads to dieback and decomposition of the felty root mat. Throw-off is also subject to erosion by wind and water. Through their burrowing activity, pikas may increase the ecosystem’s net emission of C (Qin et al. 2015), although Peng et al. (2015) could not find a direct effect of small mammals’ activity on NEE in a *Kobresia* pasture. Bare soil patches are then colonized by endemic and often monotypic rosette plants (Fig. 7K). In the long run, the windward cliffs are eroded and the open soil patches slowly develop into a gravel surface, depending on the duration and intensity of deflation (west-winds during winter, foehn from the Himalayas during the monsoon in southern Tibet; Miehe 1988). The re-establishment of *Kobresia pygmaea* in those open soils with pancake-like mats (Fig. 7G, H) is less common than the destruction of the turf, and is restricted to the eastern part of its distribution range with >300 mm/yr precipitation (Miehe and Miehe 2000).

Another common pattern of pasture degradation is found mostly on south-facing slopes, where the lower parts lack any root mat, whereas the upper slope and the ridges are covered with lawns and intact root mat. The mats form a steep cliff towards the slope with sods drifting downhill, probably along with gelifluction processes. The pattern suggests that the lower slopes had been deprived from the lawns by erosion, and the opening of the root mat may have been initiated by yak when chafing and wallowing.

Patches of dead roots covered by crustose lichens or algae (Fig. 8) are scattered across the pastures without any apparent relation to abiotic or biotic factors. The patches are rarely re-colonized by *Kobresia pygmaea*; rosettes of *Lancea tibetica* Hook.f. & Thoms. or *Kobresia macrantha* Boeckeler are more common. The most plausible explanation is dieback as a natural process with the ageing of a *Kobresia* clone, but this remains to be confirmed. Comparing the C cycle of closed lawns and crust-covered root mats by ^13^C-labeled amino acids revealed that more ^13^C remained in soil under crusts, reflecting less complete decomposition of exudates and lower root uptake (Unteregelsbacher et al. 2012). The crust patches decrease the rates of medium-term C turnover in response to the much lower C input. Very high ^13^C amounts recovered in plants from non-crust areas, and a two times lower uptake by roots under crusts indicate that very dense roots are efficient competitors with microorganisms for soluble organics. In conclusion, the altered C cycle of the crust-covered root layer is associated with strongly decreased C input and reduced medium-term C turnover.

**FIG. 8.**
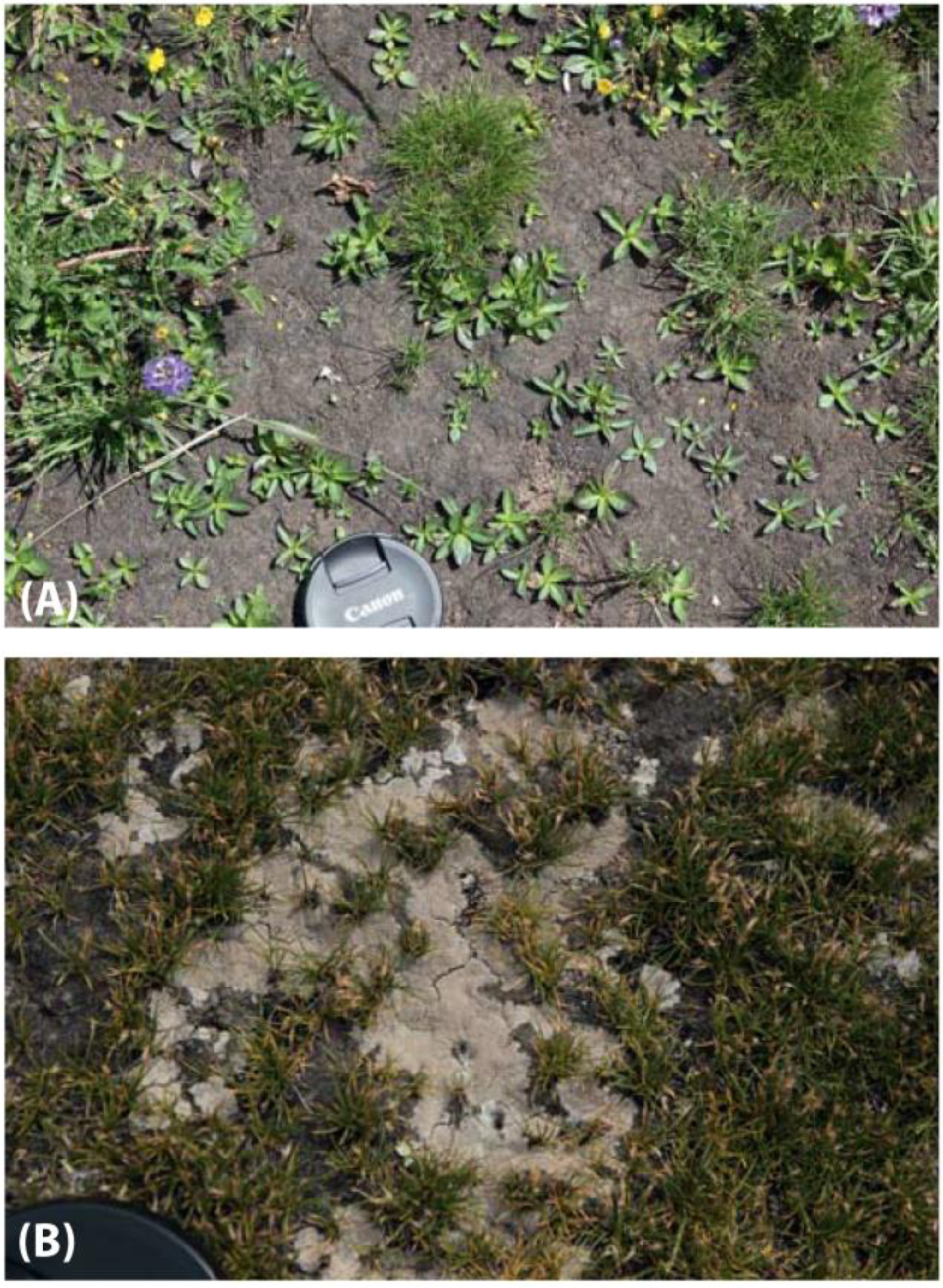
(A) Lichens and (B) Algae seal the felty *Kobresia* root mat. Photos G. Miehe 2009, 2015.

In most parts of Tibet, severe changes in plant species composition and soil fertility are spatially restricted around camps where livestock rest during night and trample frequently (i.e., piospheres; Wang et al. 2017). Heavy mechanical disturbance, in combination with excessive nutrient input (dung, urine), results in strongly altered vegetation and dieback of *Kobresia pygmaea*. The most extreme stage of degradation is a complete removal of mats (Ma et al. 1999), which can be found at landscape-scale on silty and sandy soils of the north-eastern highlands (e.g., Madoi 4300 m, 34°55’N/98°13’E; Ruoergai Plateau 3450 m, 33°35’N/102°58’E). The formation of this so-called “black soil” is widely attributed to unsustainable rangeland policies of overstocking in the late 1970 to 1980s; in some regions (Juizhi County, northeastern highlands, Li et al. 2017) in the course of fencing and privatization between 1984 and 1994. The bare soils are colonized by poisonous plants (*Aconitum luteum* H.Lév. and Vaniot) or by tiny, biennial, aromatic plants (e.g., *Artemisia* spp., *Smelowskia tibetica* Lipsky). It is here that pika densities are highest (Miehe et al. 2011*c*). In the Serxu County of the eastern highlands, 30% of the pastures have up to 4500 pika burrows/ha with 450 individuals, and an estimated harvest loss of 50% of the annual forage production (Zhou et al. 2005). Grazing exclosure experiments at Kema showed that pikas take the opportunity to avoid forage competition with livestock by excavating their dwelling burrows at a perennial grazing site or inside an undisturbed fence. Not only was the density of pikas higher inside the exclosure plots, but it was shown (by using color marking) that a large number of pikas with burrows outside the fence used its interior for foraging (at least 15%; M. Holzapfel [*personal communication*]).

Sun et al. (2015) studied the impact of pikas around Dawu (3700 m, 34°37’N/100°28’E): at highest densities of 200–300 animal/ha, these caused a decrease in species richness, vegetation cover, plant height and seasonal biomass. This pattern does, however, not seem to be the rule in the highlands because the most common disturbance indicators are forbs, and it is generally stated that pikas’ presence increases habitat diversity and plant species richness (Smith and Foggin 1999, Miehe and Miehe 2000, Smith et al. 2006), and better water infiltration and reduced erosional effects on slopes during torrential summer rains (Wilson and Smith 2015). Herders explain high pika densities as a consequence of overstocking, and not as the cause of pasture degradation (Pech et al. 2007). Pikas have been regarded as “pests” and poisoned; meanwhile the negative and long-lasting negative effects of poisoning on natural predators have been recognized and eradication programs stopped (Pech et al. 2007).

## HAVE GRAZING LAWNS FORMED AS A CONSEQUENCE OF PASTORALISM?

The ecosystem’s high share of endemics (Wu et al. 1994–2011, Miehe et al. 2011*c*) may indicate naturalness, and indeed *Kobresia* mats have been described as natural (e.g., Ni 2000, Song et al. 2004, Herzschuh and Birks 2010). With their tiny leaves, a root : shoot ratio >20, very low shoot biomass but a very large root system (Kaiser et al. 2008) and associated large C-storage, the *Kobresia pygmaea* lawns are one the world’s most extensive ecosystems with a very high below-ground share of biomass. Similar root : shoot ratios are known from other cold-adapted ecosystems including arctic tundra (Bazilevich and Tishkov 1997) and high alpine communities (Körner 2003), or from vegetation types exposed to extreme nutrient shortage like the Kwongan of western Australia (Lambers et al. 2010). Climatic parameters (Lehnert et al. 2015, 2016) including soil temperatures (Miehe et al. 2015), and the nutrient status (see above), can, however, not fully explain the prevailing structures, at least for the montane pastures. The most likely explanation of the allocation patterns found in *Kobresia* lawns is nutrient shortage in combination with intensive grazing.

The *Kobresia*-dominated pastures are commonly known as ‘alpine meadows’, which is misleading in two ways. (1) ‘meadow’ in a European sense is an agriculturally managed grassland regularly mown for livestock forage (UNESCO classification; Ellenberg and Mueller-Dombois 1965–1966). The term ‘meadow steppe’ is widely used in, for example, Mongolia (Hilbig 1995) and also refers to rangelands, yet of very different structure. In the Tibetan case the designation ‘pasture’ would be in most areas be correct (even where animal husbandry has no major impact). (2) whereas ‘alpine’ is defined strictly as a mountain climate not warm enough to allow for tree growth (Körner 2012), many ‘alpine meadows’ occur on the same slope together with isolated tree-groves (Miehe et al. 2008*c*; Fig. 9), and thus under a climate clearly suitable for tree growth. Thus pastures within the drought-line of tree growth (200–250 mm annual rainfall; Miehe et al. 2008*a*), and at altitudes below the upper tree line (3600–4800 m across the eastern highlands between the Qilian Shan and southern Tibet), have presumably replaced forests (see Fig. 1: ‘forest relics’).

**FIG. 9.**
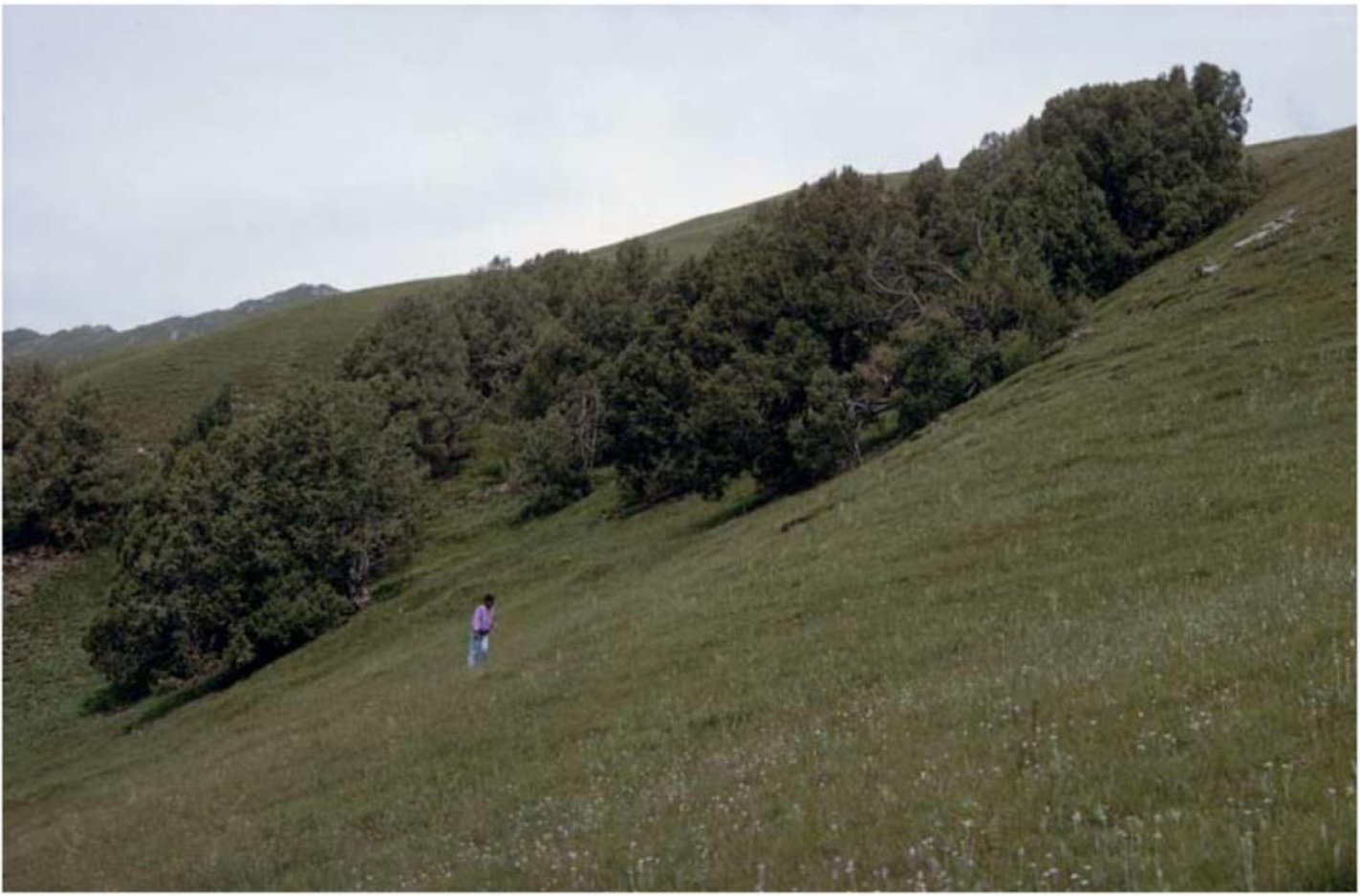
*Juniperus przewalskii* groves surrounded by *Kobresia* pastures (‘alpine meadow’, Qinghai Province, 3800 m, 35°45’N/100°14’E). Photo G. Miehe 1998.

If all pastures were natural, exclusion of livestock would have no effect. Plots of *Kobresia* grazing lawns being 2–5 cm in height were fenced in 1997 in southern Tibet (Reting; Fig. 10A), and in 2002 in the northeastern rangelands (Xinghai; Fig. 10B), both situated 300–400 m below the upper tree line. In the course of one year the lawns changed to tall Poaceae-dominated grassland 30–50 cm in height, while poisonous herbs decreased in cover (Miehe et al. 2014). The cover of *Kobresia* also decreased, but the sedge still survived under the taller grasses. Thus grazing exclusion triggered a change from a sedge-dominated mat with dwarf dicot rosettes, to a richly structured grassland with elongated dwarf sedges and rosettes in its undergrowth. In truly alpine environments (Kema), effects of fencing (erected between 2009 and 2013) were small, with grasses increasing in cover from ∼4% to 14% and reaching 15 cm in height, though this is still equivalent to a 7-fold increase of vegetation height. By contrast, *Kobresia pygmaea* leaf size increased only from ∼1.8 cm to 2.4 cm (E. Seeber [*personal communication*]). Shorter exclusion experiments in various locations in the southern highlands (Yan and Lu 2015) revealed an increase in vegetation cover, plant height and above-ground biomass, but surprisingly no change in species composition. Grazing exclusion in winter pastures (i.e. the absence of livestock interference during the vegetation period) of the northeastern ecotone towards the Alpine Steppe revealed little changes except for a decrease of the dominant grazing indicators (Harris et al. 2015). Grazing exclusion experiments thus proved that a large share of the present ‘alpine meadows’ depend for their existence on grazing.

**FIG. 10.**
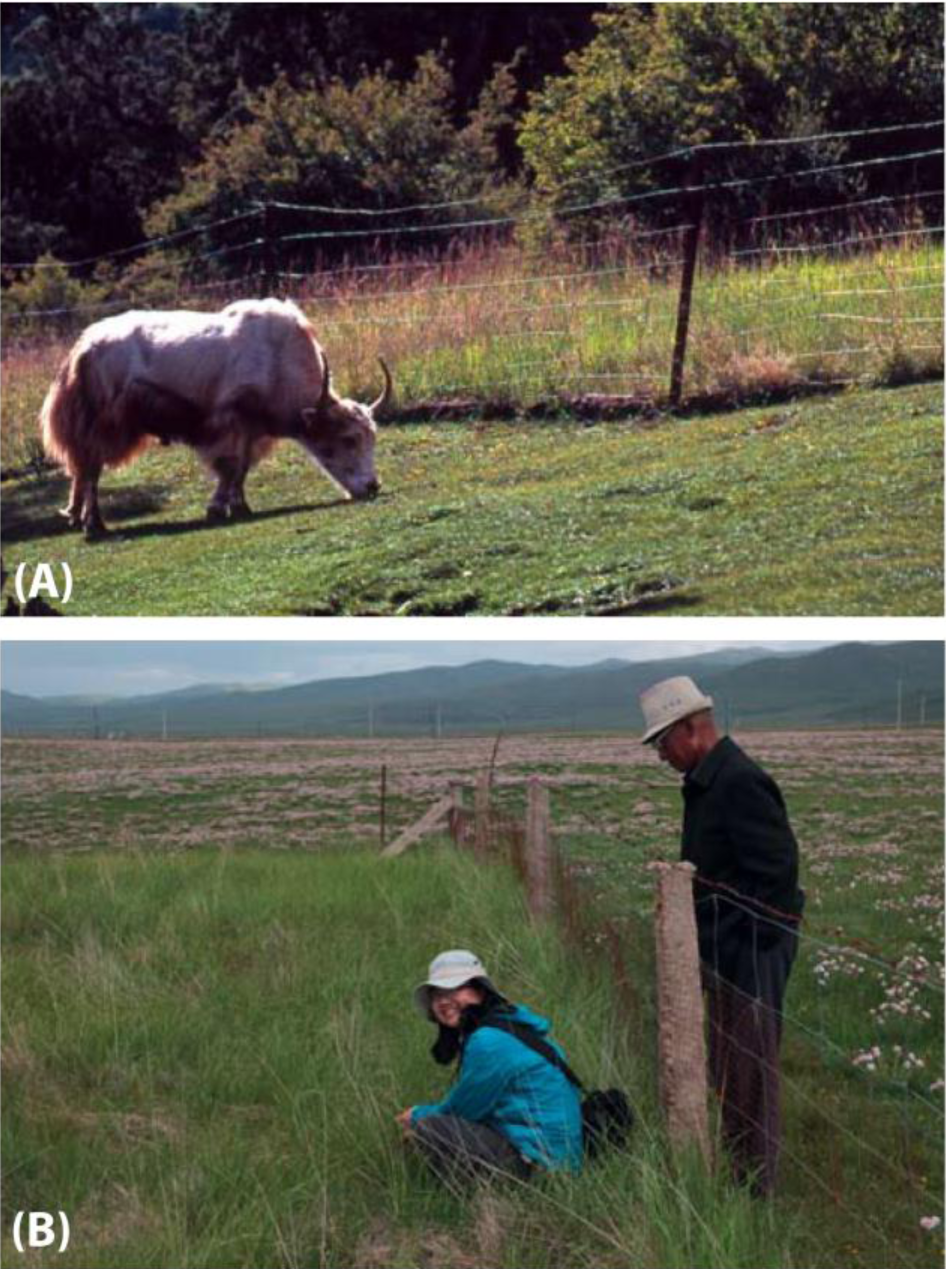
After exclusion of grazing, tall grassland overgrows lawns of *Kobresia pygmaea* giving evidence of its livestock-grazing-dependent status. (A) In Reting Forest (4250 m, 30°18’N/91°31’E) grazing lawns in a *Juniperus tibetica* woodland were fenced in 1997 and developed into a tall grassland in the first season after exclosure. Photo G. Miehe 2010. (B) In montane *Kobresia* pastures (Xinghai, 3440 m, 35°32’N/99°51’E) with a high abundance of poisonous grazing weeds (in flower: *Stellera chamaejasme*), tall *Stipa-Leymus* grasslands replaced the grazing lawns after fencing (2002) in the first season, and the grazing weeds disappeared. Photo G. Miehe 2015.

Changes from forest to grassland during the Holocene are well-documented (e.g., Herzschuh et al. 2006, Ren 2007, Miehe et al. 2014), yet the explanations are debated. As the forest decline in various areas of the highlands took place during the mid-Holocene climatic optimum (Zhang et al. 2011), a climatic driver is not plausible. Given the huge climatic niche covered by current *Kobresia* pastures, grazing offers a more parsimonious explanation. With a total height of 2–4 cm and hardly any biomass within reach of grazers (Miehe et al. 2008*b*), grazing-adapted plants like *Kobresia pygmaea* and associated species will spread at the cost of taller plants. The high root : shoot ratios and the formation of a felty turf can also be viewed in this context.

Grazing selection may also have promoted the expansion of grazing weeds. Phylogeographic studies on *Stellera chamaejasme* L. (Zhang et al. 2010), the prevalent grazing weed of *Kobresia* pastures (Fig. 10B), revealed a single haplotype over the whole of the highlands. Its presumably rapid expansion should also be explained in the context of livestock expansion.

Indeed, the Tibetan highlands have been grazed over evolutionary timescales by large herbivores (as testified by their richness, including endemic taxa; Schaller 1998). Former distribution, and natural densities, of these herbivores are largely unknown in Tibet, as they are in any other rangelands of the world. Moreover, limitations in the identification of pollen types (Miehe et al. 2009) render mapping of the historical extent of natural rangeland systems impossible. In any case animal husbandry has progressively replaced wild grazing, and there is anthracological and palynological evidence that herders extended the rangelands well into the montane forest belt (Kaiser et al. 2007, Miehe et al. 2008*a*, 2008*c*). Similar patterns have been described for the European Alps (Kral 1979) and the Himalayas (Miehe et al. 2015). This would explain why forests disappeared in spite of relatively favorable conditions, and also why montane pastures of *Kobresia pygmaea* change quickly after grazing exclusion. With increasing grazing pressure, photo assimilates are increasingly allocated below-ground. Most probably the grazing lawns, and the felty root layer, can be interpreted as an adaptation to high grazing pressure including trampling. It remains unknown if the absence of a specific endemic fauna of soil beetles can be seen as an effect of an ecosystem under stress. Due to the specific traits of the prevailing species (especially growth habit), this equilibrium system is less threatened by overstocking than is the case in other equilibrium grasslands.

## THE TIBETAN ANTHROPOCENE: FOR HOW LONG HAVE HUMANS SHAPED THIS ENVIRONMENT?

The age and intensity of the human impact on Tibetan ecosystems is debated, and estimates based on evidence from various disciplines diverge by more than 20000 years. The earliest migrations and adaptation of Tibetans to high altitude hypoxia was dated to 30 ka BP with a second migration between 10 and 7 ka BP (Qi et al. 2013), to pre-LGM and 15 ka BP (Qin et al. 2010), or to 25 ka BP (Zhao et al. 2009), yet it remains uncertain where this mutation occurred (Madsen 2016). Human population genomic data suggest that the most critical *EPAS1* genetic variant for hypoxia adaptation of Tibetans derived from extinct Denisovan people who hybridized with ancestors of Tibetans and Han Chinese (Huerta-Sánchez et al. 2014). Whereas the Tibetan population retained this genetic variant, and had its frequency increased due to the strong natural selection, it was lost in the Han Chinese and other groups. The widespread occurrence of artefacts, including stone tools, provide archeological evidence for human presence on the plateau (Brantingham and Gao 2006, Bellezza 2008, Chen et al. 2015), but the dating of scattered surface remains difficult. While archeology-based estimates of the time for the first intrusion of hunters range between 30 and 8 ka BP (Aldenderfer 2006, Brantingham et al. 2007), ^14^C- and OSL-dated remains suggest a more recent date suggesting that hunting parties first travelled in the region between 16 and 8 ka BP. The relevance of hand – and footprints in tufa-sediments north of Lhasa (Chusang) remains obscure as data range from 26 ka (Zhang and Li 2002) to 7.4 ka BP (Meyer et al. 2017). Obsidian tools dated between 9.9 and 6.4 ka BP have been transported over more than 900 km (Perreault et al. 2016). In any case, humans have most probably travelled within the region during the Last Glacial Maximum (LGM). Long-term residential groups of hunters, or perhaps early pastoralists, may have settled in the area between 8 and 5 ka BP (Madsen et al. 2017), a date supported by independent evidence from the genomic signature of yak domestication (Qiu et al. 2015).

So far direct proof of early human impact on vegetation structures has not emerged. The presence of humans is commonly associated with the use of fire, wherever fuel loads in the dry season are high enough for lighting (Bond and Keeley 2005). Fire traces observed in Tibetan sediments may relate to human action (see Fig. 11). As highland plants lack obvious adaptations to fire (e.g., no pyrophytes as present in the Boreal Forest, the South African Fynbos, or the Australian Kwongan) it seems unlikely that fire had been present over evolutionary time-scales prior to human arrival. Lightning occurs nearly exclusively during the rainy season, followed by torrential rains. Between 2002 and 2005, 96% of lightning records of 36 climate stations in Qinghai Province occurred during the rainy season (Meteorological Service Qinghai [*personal communication*]); lightning imaging sensors gave similar results for the central and eastern highlands (Qie et al. 2003).

**FIG. 11.**
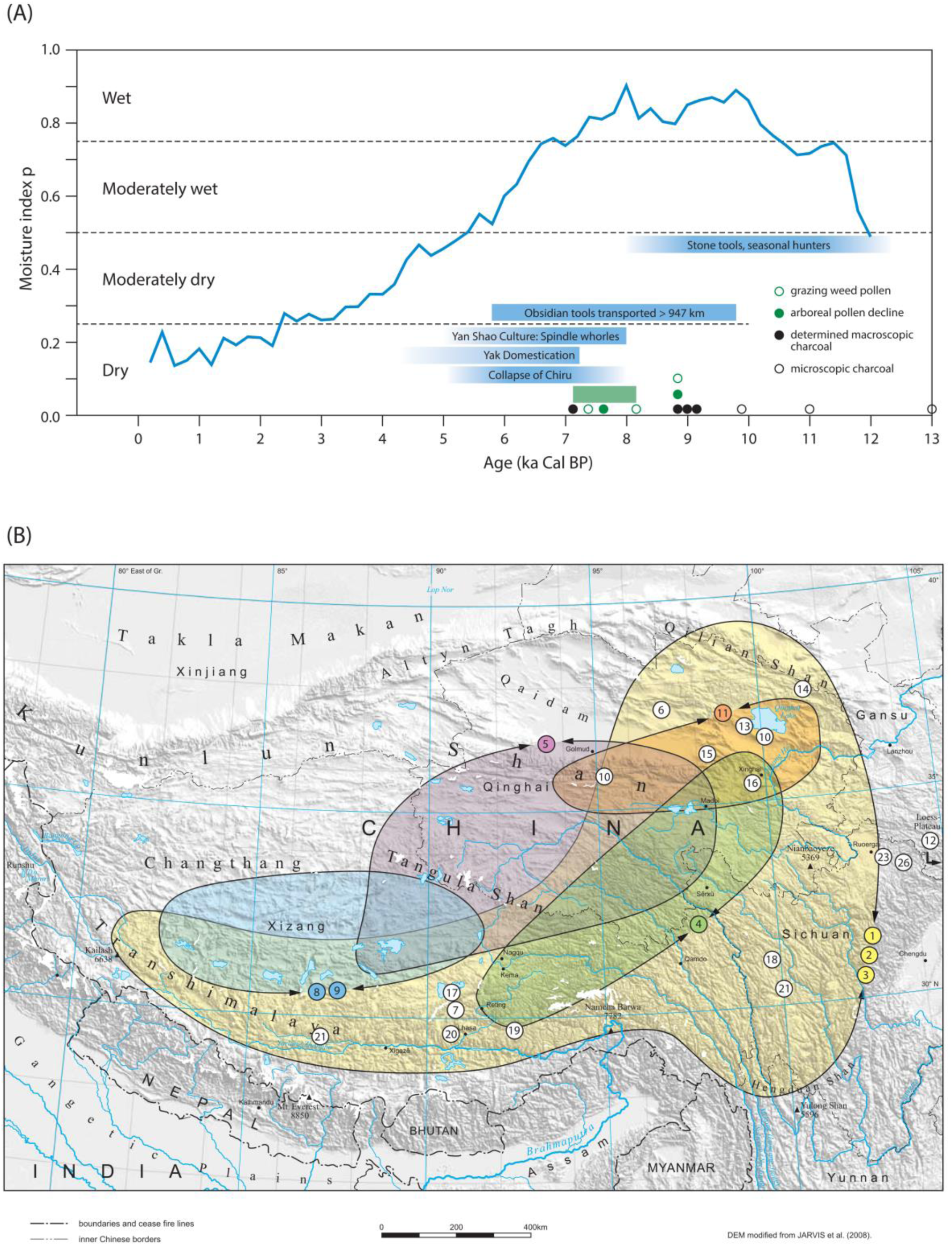
Summary diagram of moisture conditions and evidence of human impact during the early Anthropocene in the Tibetan highlands: The use of fire by early hunters changed vegetation structures from forest to grassland from the time of the mid Holocene climatic optimum. The present grazing lawns have developed with increasing grazing pressure since the onset of pastoralism. After data of Zhang et al. 2011, and the sources mentioned in table 2 (all ages in ka cal. BP). References are given in the Supplementary Material Table S2.

The seasonality of the highland climate would favors the use of fire as a tool to modify vegetation structures. This is especially true for the mid-Holocene climatic optimum between 10 and 5 ka BP, when summer growing conditions were wetter and warmer than present (Zhang et al. 2011), resulting in presumably high fuel load available for burning during the dry cold winter. Fires lit by hunters were most probably the first impact, as they are associated with the first intrusion of humans in other parts of the world (e.g. Ogden et al. 1998). Landscape-scale management by burning has probably intensified with the introduction of livestock grazing: forests were burnt to provide better pastures, and to destroy places of concealment for predators. Charcoal of *Picea* (*P. crassifolia* Komarov) and *Juniperus* (*J. przewalskii* Komarov) has been found in the north-eastern highland pastures where current precipitation is double the minimum amount of rainfall necessary for tree growth. Moreover, these sites are situated at altitudes 200–400 m below the upper tree line (Miehe et al. 2008*c*). Dating of the charcoal implies ages between 10.0 and 7.4 ka BP (Kaiser et al. 2007). On the south-eastern slope (Hengduan Shan, 4200 m, 31°06’N/99°45’E), fire has been recorded since 13 ka BP, and a decline of *Betula* pollen between 8.1 and 7.2 ka BP occurs synchronously with charcoal peaks (Kramer et al. 2010). Other charcoal records of burned shrubs date back to 48.6 ka BP (Kaiser et al. 2009*b*). Phylogeographic analyses of now disjunct forest relicts (*Picea crassifolia*, *Juniperus przewalskii*) with shared haplotypes across the northeastern pastures suggests that post-glacial recolonization resulted in continuous forests (Zhang et al. 2005, Meng et al. 2007), that have been opened up and fragmented more recently by pasturing.

**TABLE 2.**
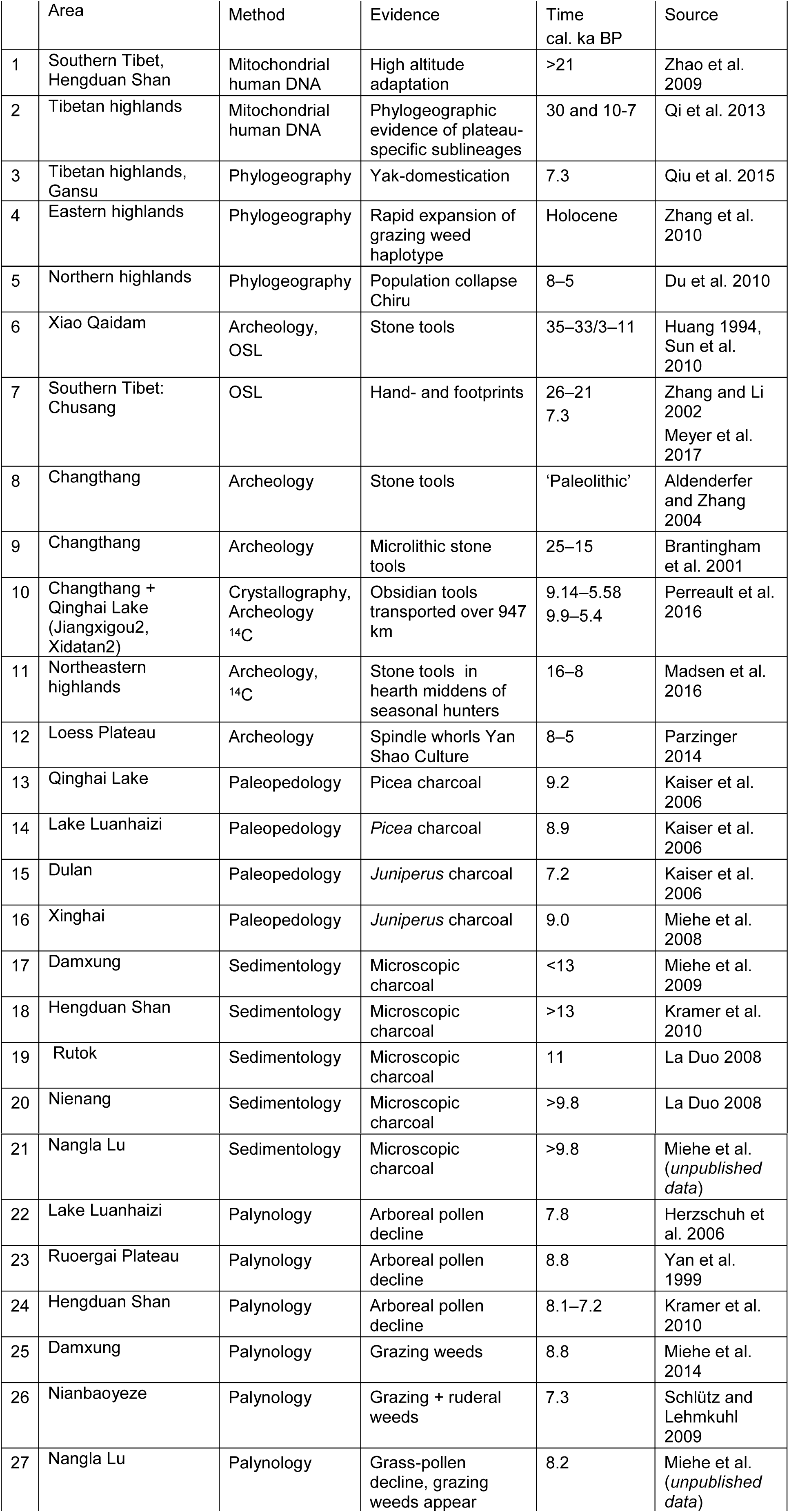
Summary table for fig. 11; references are given in the Supplementary Material Table S2.

|  | Area | Method | Evidence | Time cal. ka BP | Source |
| --- | --- | --- | --- | --- | --- |
| 1 | Southern Tibet, Hengduan Shan | Mitochondrial human DNA | High altitude adaptation | >21 | Zhao et al. 2009 |
| 2 | Tibetan highlands | Mitochondrial human DNA | Phylogeographic evidence of plateau-specific sublineages | 30 and 10-7 | Qi et al. 2013 |
| 3 | Tibetan highlands, Gansu | Phylogeography | Yak-domestication | 7.3 | Qiu et al. 2015 |
| 4 | Eastern highlands | Phylogeography | Rapid expansion of grazing weed haplotype | Holocene | Zhang et al. 2010 |
| 5 | Northern highlands | Phylogeography | Population collapse Chiru | 8–5 | Du et al. 2010 |
| 6 | Xiao Qaidam | Archeology, OSL | Stone tools | 35–33/3–11 | Huang 1994, Sun et al. 2010 |
| 7 | Southern Tibet: Chusang | OSL | Hand- and footprints | 26–21<br>7.3 | Zhang and Li 2002<br>Meyer et al. 2017 |
| 8 | Changthang | Archeology | Stone tools | 'Paleolithic' | Aldenderfer and Zhang 2004 |
| 9 | Changthang | Archeology | Microlithic stone tools | 25–15 | Brantingham et al. 2001 |
| 10 | Changthang + Qinghai Lake (Jiangxigou2, Xidatan2) | Crystallography, Archeology <sup>14</sup>C | Obsidian tools transported over 947 km | 9.14–5.58<br>9.9–5.4 | Perreault et al. 2016 |
| 11 | Northeastern highlands | Archeology, <sup>14</sup>C | Stone tools in hearth middens of seasonal hunters | 16–8 | Madsen et al. 2016 |
| 12 | Loess Plateau | Archeology | Spindle whorls Yan Shao Culture | 8–5 | Parzinger 2014 |
| 13 | Qinghai Lake | Paleopedology | *Picea* charcoal | 9.2 | Kaiser et al. 2006 |
| 14 | Lake Luanhaizi | Paleopedology | *Picea* charcoal | 8.9 | Kaiser et al. 2006 |
| 15 | Dulan | Paleopedology | *Juniperus* charcoal | 7.2 | Kaiser et al. 2006 |
| 16 | Xinghai | Paleopedology | *Juniperus* charcoal | 9.0 | Miehe et al. 2008 |
| 17 | Damxung | Sedimentology | Microscopic charcoal | <13 | Miehe et al. 2009 |
| 18 | Hengduan Shan | Sedimentology | Microscopic charcoal | >13 | Kramer et al. 2010 |
| 19 | Rutok | Sedimentology | Microscopic charcoal | 11 | La Duo 2008 |
| 20 | Nienang | Sedimentology | Microscopic charcoal | >9.8 | La Duo 2008 |
| 21 | Nangla Lu | Sedimentology | Microscopic charcoal | >9.8 | Miehe et al. (*unpublished data*) |
| 22 | Lake Luanhaizi | Palynology | Arboreal pollen decline | 7.8 | Herzschuh et al. 2006 |
| 23 | Ruoergai Plateau | Palynology | Arboreal pollen decline | 8.8 | Yan et al. 1999 |
| 24 | Hengduan Shan | Palynology | Arboreal pollen decline | 8.1–7.2 | Kramer et al. 2010 |
| 25 | Damxung | Palynology | Grazing weeds | 8.8 | Miehe et al. 2014 |
| 26 | Nianbaoyeze | Palynology | Grazing + ruderal weeds | 7.3 | Schlütz and Lehmkuhl 2009 |
| 27 | Nangla Lu | Palynology | Grass-pollen decline, grazing weeds appear | 8.2 | Miehe et al. (*unpublished data*) |

Charcoal and pollen records point to a forest decline (*Picea*, *Betula*) after between 8 and 6 ka BP (Shen et al. 2005, Herzschuh et al. 2006, Cheng et al. 2010, Miehe et al. 2014), at a period that, according to the human impact-independent proxy of Ostracod assemblages (Mischke et al. 2005), was the most favorable climatic period of the Holocene. As the pollen record does not show the usual post-fire succession (*Epilobium>Populus>Betula*), it is plausible that humans suppressed forest recovery because they preferred grassland for their livestock instead of forest. In the forest ecotone of the eastern highland slope (Nianbaoyeze, 33°22’N/101°02’E), the fire record covers the whole of the Holocene, while pollen of ruderals (*Tribulus*) and grazing weeds (*Stellera chamaejasme*) start to appear around 7.3 ka BP (Schlütz and Lehmkuhl 2009, Miehe et al. 2014). Pollen-diagrams from *Kobresia*-swamps of the Ruoergai Plateau (i.e. Zoige Basin) (650 to 780 mm/yr; Yan et al. 1999) show an abrupt forest pollen decline around 8.8 ka BP, followed by increased values of Poaceae and Chenopodiaceae, similar to a pollen-sequence typical for the ‘landnam’ in Europe (Frenzel 1994).

In southern Tibet (Damxung 4250 m, 30°22’N/90°54’E; Miehe et al. 2009), the charcoal record since the Late Glacial is uninterrupted; and the most common human-indicator pollen types (*Stellera chamaejasme*, *Pterocephalus*, *Cyananthus*, *Plantago*) first appear around 8.5 ka BP. As the endemic large herbivores in Tibet have similarly selective grazing habits to those of livestock (Miller and Schaller 1996), the increase of grazing weeds could well be a result of increased numbers of wild herbivores. Phylogeographical analysis of *Panthelops hodgsonii* Abel (chiru, Tibet antelope), however, shows a clear decrease in herd-size during this period (Du et al. 2010), probably synchronous with the domestication of the yak (7.3 ka BP; Qiu et al. 2015). Spores of dung fungi (*Sporormiella*), indicate pasture degradation in Damxung over the last 1200 years, and *Glomus* spores show that roots had been exposed through erosion. Other sites in southern Tibet (Rutok, 29°41’N/92°16’E, Nienang, 29°43’N/90°42’E; van Leeuwen in La Duo 2008) show charcoal peaks between 11.3 and 6 ka BP.

The archeo-zoological record of early animal husbandry and agriculture is still very limited; earliest fossil records of domestic yak reveal ages of only 3.8 ka BP (Flad et al. 2007), which are similar to the oldest records of cereals (barley in the northeastern highlands: 4 ka BP, in the Yarlung Zhangbo valley: 3.4 ka BP; *Setaria italica* (L.) P.Beauv. on the eastern slope of the highlands: 4.6 ka BP; Chen et al. 2015, d’ Alpoim Guedes 2015). The permanent human occupation of the highlands is thought to have depended on mutual exchange between cereal farmers (trading barley as the key staple food of Tibetans) and pastoralists (giving animal products and salt), and has thus been dated as younger than 3.6 ka BP (Chen et al. 2015). Recent genomic evidence, however, implies a much older date for the domestication of the yak (7.3 ka BP; Qiu et al. 2015).

One of the most important prehistoric excavations in China, near Xian on the Loess Plateau, is possibly also relevant for the question of the onset of pastoralism in the Tibetan highlands: In settlements of the Yangshao Culture (7–5 ka BP; Parzinger 2014), spindle whorls, found in large quantities, most probably testify to the weaving of sheep wool. Whether similarly aged bones of Caprini found at those sites are of domestic origin or not, is unclear (Flad et al. 2007). Given that sheep must have been introduced from their center of domestication in the mountains of the Middle East (Zeder and Hesse 2000), and that Tibet is 1600 km closer to this center of origin than is Xian, pastoralism may have started earlier in the former than the latter. If we hypothetically apply the same rate of diffusion of the Neolithic Package from the center of domestication towards Europe (3 km/yr from the Zagros Mountains in western Iran to the south-east European Vojvodina; Roberts 1998), domestic sheep and goats may have reached the Tibetan highlands around 8.6 ka BP.

There thus is evidence from various sources indicating that human land use has established shortly after the end of the glacial period. The question of when increased livestock numbers first had an impact on plant cover, and on the below-ground C allocation that forms the felty root mats, is still, however, unanswered.

## CONCLUSIONS

1. The *Kobresia* pastures of the Tibetan highlands are a nutrient- and water-limited high-altitude ecosystem. They form equilibrium rangeland systems, yet their vulnerability to grazing degradation is limited due to the prevalence of a dwarf sedge with its main above-ground phytomass below the grazing reach of livestock.
2. *Kobresia* mats are characterized by a felty root layer that represents very large carbon (C) store. Natural degradation phenomena with polygonal crack patterns and the drifting apart of polygonal sods are, however, widespread. The widening of the cracks is accompanied by high SOC losses (∼5 kg C/m^2^). Overall up to 70% of the SOC stock was lost in comparison with intact swards of alpine *Kobresia* pastures. Assuming that the whole *Kobresia* ecosystem has suffered to a similar extent, this would imply a total SOC loss of 0.6 Pg C for the whole southeastern highlands. Consequently high amounts of C are released back to the atmosphere as CO_2_, or are deposited in depressions and rivers.
3. Pastoralism may have promoted dominance of *Kobresia pygmaea* and is a major driver for below-ground C allocation and C sequestration, stabilizing these root mats with their distinctive C allocation patterns. Grazing exclosure experiments show that the larger below-ground C allocation of plants, the larger amount of recently assimilated C remaining in the soil, and the lower soil organic-matter derived CO_2_ efflux create a positive effect of moderate grazing on soil C input and C sequestration in the whole ecosystem.
4. Due to the highlands’ relevance for atmospheric circulation patterns, surface properties of these pastures have an impact on large and possibly global spatial scales. The removal of the lawns, caused by climatic stress as well as excessive human impact leads to a shift from transpiration to evaporation in the water budget, followed by an earlier onset of precipitation and decreasing incoming solar radiation, resulting in changes in surface temperature, which feedback on changes in atmospheric circulations on a local to regional scale.
5. The age of the world’s largest alpine ecosystem, and its set of endemic plants and animals, remains a matter of considerable dispute, though the degree of this uncertainty is rarely admitted (reviews in Liu et al. 2014, Favre et al. 2015, Schmidt et al. 2015, Renner 2016). Further calibrations based on genomic data and fossils (where available) are needed to clarify evolutionary relationships and divergences between the currently recognized species of *Kobresia*.
6. The paleo-environmental evidence, as well as simulations, suggests that the present grazing lawns of *Kobresia pygmaea* are a synanthropic ecosystem that developed through selective free-range grazing of livestock. The age of the present grazing lawns, however, is not yet known. The presence of humans using fire and replacing forests by grassland may date back as far as the LGM, while archeological evidence for such an early onset of pastoralism is missing. A multi-proxy approach, however, suggests a mid-Holocene climatic optimum age.
7. The traditional migratory, and obviously sustainable, rangeland management system conserved and increased the C stocks in the turf and its functioning in the regional and global C cycles. However, rangeland management decisions within the past 50 years have caused widespread overgrazing leading to erosion and reducing the C sink strength. Considering the large area of the grasslands, even small reductions in C sequestration rate would affect the regional C balance, with possible impacts on the future climate of China and beyond.

## ACKNOWLEDGEMENTS

Fieldwork was carried out with support of the German Research Council (DFG SPP 1372); the support of its steering committee is especially acknowledged. The research station at Kema was established with funding of the VW Foundation within the framework of the Marburg – Lhasa University Partnership Program. Assessments on pasture degradation have been supported by the German Federal Government’s Ministry of Education and Research (BMBF-CAME framework). The work of Jianquan Liu was supported by the Ministry of Science and Technology of the People’s Republic of China (2010DFA34610), International Collaboration 111 Projects of China.

Special thanks go to Christiane Enderle for preparing the figures and maps. We further thank Dawa Norbu, Eva Falge, Eugen Görzen, Gwendolin Heberling, Hanna Meyer, Jürgen Leonbacher, Klaus Schützenmeister, Lang Zhang, Lena Becker, Lobsang Dorji, Olga Shibistova, Sabrina Träger, Stefan Pinkert, Thomas Leipold and Yue Sun for their support during field work, and local people and landowners for their hospitality and cooperation.

